# Zα domain-dependent ZBP1 condensate formation induces an amyloidal necroptotic signalling complex

**DOI:** 10.1101/2025.06.06.658343

**Authors:** Josephine Nemegeer, Aynur Sönmez, Evelien Dierick, Katrien Staes, Leslie Naesens, Peter Vandenabeele, Denis L.J. Lafontaine, Jonathan Maelfait

## Abstract

ZBP1 restricts viral replication by inducing host cell death. ZBP1 recognises Z-RNA or Z-DNA, left-handed double-stranded RNA or DNA structures that accumulate after virus infection. How the interaction between Z-RNA/DNA and ZBP1 governs its activation and how this mediates downstream signalling remains unclear. Using herpes simplex virus 1 (HSV-1) as an activator of human ZBP1 we find that binding of the N-terminal Zα domains of ZBP1 to Z-RNA induces ZBP1 condensate formation. This then mediates oligomerisation of the RIP homotypic interaction motifs (RHIMs) of ZBP1 establishing an amyloidal signalling complex with RIPK1 and RIPK3 that induces necroptotic cell death. We find that the kinase activity of RIPK1 is essential for RIPK1 and RIPK3 oligomerisation downstream of human ZBP1. Finally, the HSV-1-encoded RHIM-containing protein ICP6, does not interfere with Zα domain-mediated ZBP1 condensate formation, but instead prevents downstream RIPK1 and RIPK3 oligomerisation thereby inhibiting necroptosis and promoting viral growth. Together, this shows that ZBP1 condensate formation restricts HSV-1 infection by promoting host cell necroptosis.

## Introduction

Recognition of foreign or self-nucleic acids by nucleic acid-sensing pattern recognition receptors activates an innate antiviral immune response (Bartok & Hartmann, 2020; Tan *et al*, 2018). The nucleic acid sensor Z-DNA binding protein 1 (ZBP1), previously referred to as DAI (Takaoka *et al*, 2007), is activated by a wide range of stimuli including DNA and RNA viruses, chemotherapeutics, and genetic mutations (DeAntoneo *et al*, 2023; Karki & Kanneganti, 2023; Maelfait & Rehwinkel, 2023). As such, ZBP1 is involved in multiple pathophysiological processes such as antiviral defence, anticancer immunity and autoinflammation.

ZBP1 activation depends on the interaction with Z-nucleic acids, double-stranded (ds) RNA or dsDNA helices that have adopted a left-handed Z-conformation. Z-nucleic acids, including Z-RNA (Hall *et al*, 1984) and Z-DNA (Wang *et al*, 1979), are predicted to be relatively rare in healthy cells. This presumption is based on the fact that in physiological solution Z-conformations are energetically unfavourable as opposed to their respective right-handed A- and B-conformers (Krall *et al*, 2023; Rich *et al*, 1984). Proteins that bind Z-nucleic acids, including ZBP1, utilise Zα domains to stabilise Z-RNA/Z-DNA sequences either by binding to pre-existing Z-sequences and/or through active conversion of right-handed A-RNA and B-DNA into the Z-conformation (Herbert *et al*, 1997). ZBP1 contains two N-terminal Zα domains, termed Zα1 and Zα2 (see Fig. 1A). Both Zα domains bind to Z-DNA, mediate B-to-Z-DNA conversion (Deigendesch *et al*, 2006; Ha *et al*, 2008; Schwartz *et al*, 2001) and contribute to ZBP1 signalling, although the Zα2 domain has a dominant function (Amusan *et al*, 2025; Maelfait *et al*, 2017; Sridharan *et al*, 2017; Thapa *et al*, 2016).

**Figure 1.**
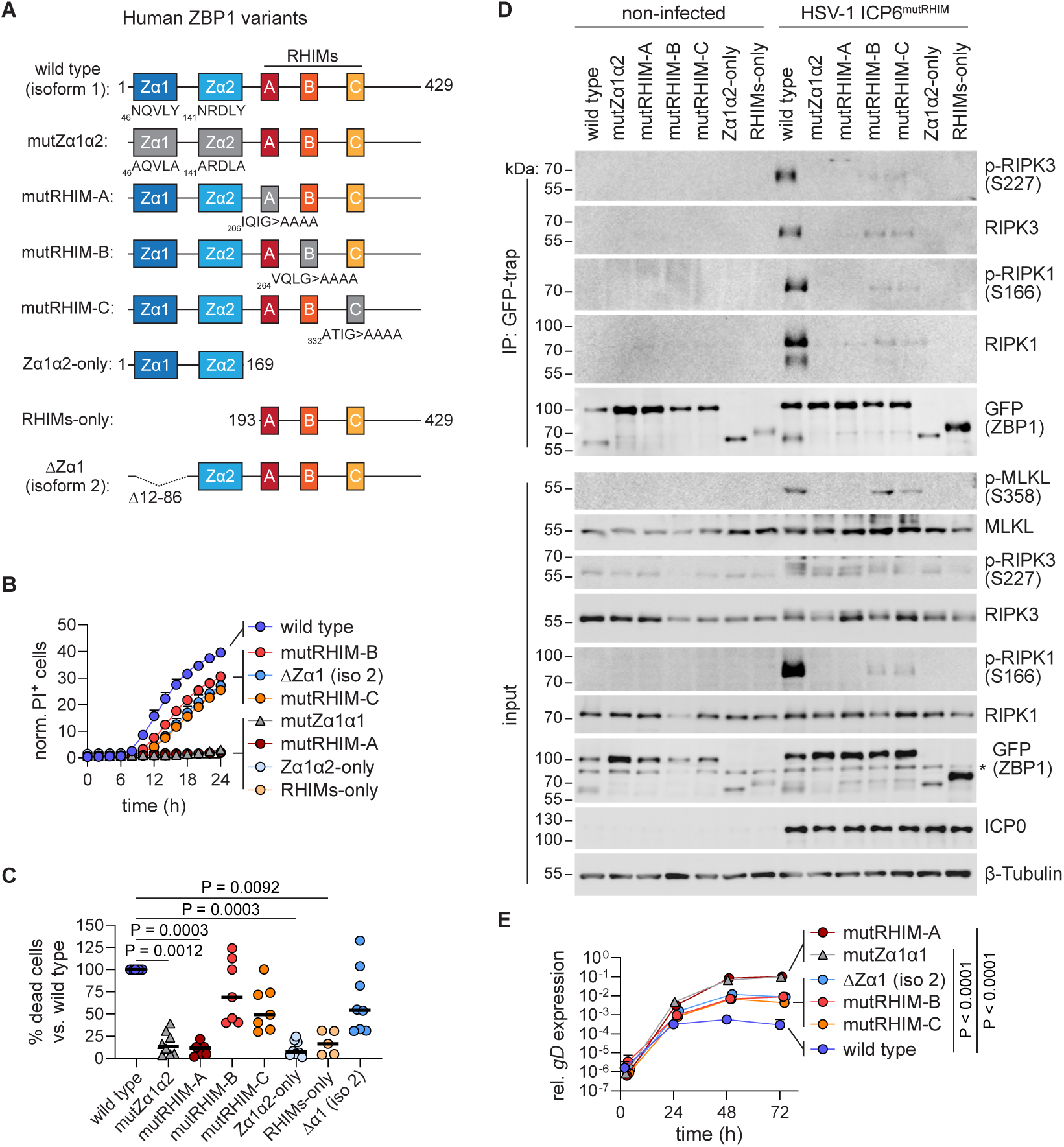
Human ZBP1-induced necroptosis depends on Zα domains and RHIMs. **(A**) Schematic overview of the different human ZBP1 variants used in this study. Wild type ZBP1 (isoform 1) contains two Zα domains (Zα1 and Zα2), followed by three RIP homotypic interaction motifs (RHIM-A, RHIM-B and RHIM-C) and a C-terminal tail. Amino acid mutations and the positions are indicated. The natural splice variant, ZBP1 isoform 2 (ΔZα1), does not contain amino acids 12-86 comprising Zα1. **(B)** HT-29 cells transduced with doxycycline-inducible lentivectors expressing the indicated C-terminally eGFP-V5 tagged human ZBP1 variants were infected with HSV-1 ICP6^mutRHIM^ at a multiplicity of infection (MOI) of 5. Unless stated otherwise, cellular assays were performed based on leaky expression of ZBP1 or its variants from the doxycycline-inducible promotor. Cell death was quantified by measuring propidium iodide (PI) uptake every 2 hours using Incucyte cell imaging. The number of PI^+^ cells per image at each time point was divided by the percentage of confluency to obtain normalised values plotted as “norm. PI^+^ cells” on the Y-axis. Lines represent a sigmoidal, 4PL fit. **(C)** Percentage of norm. PI^+^ or Sytox Green^+^ cells 18 hours after HSV-1 ICP6^mutRHIM^ infection (MOI of 5). Each data point represents an independent experiment. Values for wild type ZBP1 were set at 100 % within each experiment. P values by Kruskal-Wallis test. **(D)** HT-29 cells expressing the indicated eGFP-V5-tagged human ZBP1 variants were infected for 9 hours with HSV-1 ICP6^mutRHIM^ (MOI of 5). ZBP1-eGFP-V5 was immunoprecipitated (IP) using GFP-Trap beads and input and IP samples were analysed by western blotting. * represents a non-specific signal. **(E)** HT-29 cells expressing human ZBP1-eGFP-V5 variants were infected with HSV-1 ICP6^mutRHIM^ (MOI of 0.1). Viral replication was measured at the indicated time points by determining relative mRNA expression of the HSV-1 *gD* gene using RT-qPCR. P values by 2way ANOVA.

While engagement of ZBP1 is mediated by interactions between Z-nucleic acids and its Zα domains, signalling downstream of ZBP1 depends on the recruitment of receptor-interacting serine/threonine-protein kinase 1 (RIPK1) and RIPK3. Direct interactions between ZBP1 and either RIPK1 or RIPK3 are mediated through RIP homotypic interaction motifs (RHIMs), which are present in all three proteins (Kaiser *et al*, 2008; Rebsamen *et al*, 2009; Sun *et al*, 2002). In conditions that stimulate necroptosis, RIPK1 and RIPK3 organise into β-amyloidal fibrils composed out of two antiparallel β-sheets that are formed by stacking of the RHIMs of RIPK1 and/or RIPK3 (Li *et al*, 2012; Liu *et al*, 2024; Mompean *et al*, 2018; Wu *et al*, 2021a; Wu *et al*, 2021b). This amyloidal assembly, called the necrosome, is thought to serve as a signalling platform for RIPK3-mediated MLKL phosphorylation and oligomerisation resulting in necroptotic cell death (Chen *et al*, 2022; Cho *et al*, 2009; He *et al*, 2009; Sun *et al*, 2012; Zhang *et al*, 2009; Zhao *et al*, 2012). While RIPK1 and RIPK3 both contain a single RHIM, ZBP1 contains three RHIMs termed RHIM-A, -B and -C that differ in amino acid composition and possibly function (see Fig. 1A) (Kaiser *et al*., 2008; Rebsamen *et al*., 2009; Sun *et al*., 2002). Recruitment of RIPK1 and RIPK3 to ZBP1 activates the proinflammatory transcription factor NF-κB (Kaiser *et al*., 2008; Peng *et al*, 2022; Rebsamen *et al*., 2009), or induces caspase-8 (CASP8)-dependent apoptosis or MLKL-dependent necroptosis (Kuriakose *et al*, 2016; Thapa *et al*., 2016; Upton *et al*, 2012). It is not clear yet whether these different downstream signalling pathways originate from the same or from distinct signalling complexes. It is likely that -analogous to the TNFR1-induced signalling complex-the outcome of ZBP1 activation is determined by post-translational modifications of signalling components such as phosphorylation, K63- or M1-linked ubiquitination, or proteolytic processing of RIPK1 (Clucas & Meier, 2023; Huyghe *et al*, 2023). At least in mouse cells, necroptosis downstream of ZBP1 is kept in check through the cleavage of RIPK1 by CASP8 (Imai *et al*, 2024; Schwarzer *et al*, 2020; Yang *et al*, 2020). Of note, only human and not mouse ZBP1 has been reported to induce NF-κB activation (Koerner *et al*, 2024; Peng *et al*., 2022), suggesting that mouse and human ZBP1 signalling complexes are differentially regulated.

How the interaction of Z-nucleic acids with its Zα domains activates ZBP1 and how this coordinates downstream signalling is not clear yet. Moreover, despite some notable species differences, such as the capacity to activate NF-κB, most studies on ZBP1 signalling have been performed in mouse systems. We therefore undertook a protein domain/function mapping approach to study the molecular mechanisms that govern human ZBP1 activation using herpes simplex virus-1 (HSV-1) as an activator of ZBP1 (Guo *et al*, 2018). We find that binding of Z-RNA to ZBP1’s Zα domains induces the formation of partially dynamic ZBP1 condensates independently of the RHIMs. We propose that by increasing the local concentration of ZBP1 in these condensates, the Z-RNA-Zα domain-mediated interactions then promote proximity-induced homotypic interactions between the RHIMs of ZBP1, establishing an amyloidal ZBP1 signalling platform that recruits and activates RIPK1 and RIPK3 resulting in MLKL activation and necroptosis. Replacing the Zα domains of ZBP1 by a self-oligomerising CRY2olig domain (Taslimi *et al*, 2014), which enabled light-inducible/ligand-independent RHIM-mediated ZBP1 oligomerisation, further confirmed the capacity of the RHIMs of ZBP1 to form amyloid-like assemblies. A RHIM present in the HSV-1 immediate early protein ICP6, which has previously been described to inhibit necroptosis in human cells (Guo *et al*., 2018; Guo *et al*, 2015; Huang *et al*, 2015; Wang *et al*, 2014), inhibits RIPK1 and RIPK3 recruitment to ZBP1. In contrast, ICP6 does not prevent the formation of ZBP1 foci and only minimally affects ZBP1 amyloid formation. Finally, in contrast to its mouse orthologue (Upton *et al*., 2012), human ZBP1 induces necroptosis in a manner that strictly depends on the kinase activity of RIPK1. Mechanistically. we find that the enzymatic activity of RIPK1 is not required for ZBP1 amyloid formation, but instead promotes stable RIPK1/RIPK3 oligomerisation. Together, we show that the Z-RNA-induced formation of amyloidal ZBP1 condensates acts as a signalling platform to induce host cell necroptosis.

## Results

### Endogenous human ZBP1 induces necroptosis in response to HSV-1 infection

Others and we previously showed that ectopic expression of human ZBP1 in the human HT-29 colorectal adenocarcinoma cell line can induce either NF-κB, CASP8 or MLKL activation (de Reuver *et al*, 2022; Guo *et al*., 2018; Peng *et al*., 2022; Zhang *et al*, 2020). This indicates that HT-29 cells support the formation of inflammatory, apoptotic or necroptotic ZBP1 signalling complexes rendering this cell line suitable for studying human ZBP1 activation. Since these studies relied on overexpression we first asked whether endogenous ZBP1 is active in HT-29 cells. Human ZBP1, similar to the mouse orthologue and in line with previous work (Fu *et al*, 1999; Pham *et al*, 2006), is an interferon-stimulated gene (Fig. S1A,D,F). To control for the specificity of the antibody we included protein lysates from the MM1S multiple myeloma cell line, which constitutively expresses high levels of ZBP1 (Ponnusamy *et al*, 2022) (Fig. S1F) or lysates from HT-29 cells in which ZBP1 expression was depleted by siRNAs or CRISPR-Cas9 (Fig. S1D,E,F). Two ZBP1 species with apparent molecular weights around 45 and 55 kDa were detected in both HT-29 and MM1S cells, likely representing two *ZBP1* isoforms encoded by Ensembl transcripts ZBP1-201 or ZBP1-202 and in previous studies annotated as isoform 1/ZBP1(L) or isoform 2/ZBP1(S), respectively (Nassour *et al*, 2023; Ponnusamy *et al*., 2022). Transcript ZBP1-201, containing 8 exons, constitutes the reference sequence and codes for a protein containing two Zα domains (Zα1 and Zα2) followed by three RHIMs (termed RHIM-A, -B and -C) and a predicted disordered C-terminal tail (Fig. 1A; wild type, isoform 1). Transcript ZBP1-202 arises from exon 2 skipping and translates into a ZBP1 protein lacking the first Zα domain (Fig. 1A; ΔZα1, isoform 2) (Rothenburg *et al*, 2002). Although isoform 2 is the predominant splice variant, at least in HT-29 cells, we performed our domain/function studies on isoform 1 comprising both Zα domains and hereafter referred to as wild type ZBP1.

To determine whether endogenous ZBP1 is functionally active, we stimulated HT-29 cells with IFN-α2 to induce ZBP1 expression and subsequently infected these cells with an HSV-1 strain encoding RHIM-mutant ICP6 (HSV-1 ICP6^mutRHIM^), which is unable to inhibit ZBP1-dependent necroptosis (Guo *et al*., 2018; Huang *et al*., 2015). IFN-α2 pre-treatment sensitised HT-29 cells to HSV-1 ICP6^mutRHIM^-induced cell death in a dose-dependent manner (Fig. S1B). siRNA- or CRISPR-Cas9-mediated depletion of ZBP1 expression prevented cell death after HSV-1 ICP6^mutRHIM^ infection, while TNF receptor 1 (TNFR1)-induced necroptosis, after stimulation with TNF, the SMAC inhibitor BV6 and the pan-caspase inhibitor zVAD-fmk (TNF/BV6/zVAD) was unaffected by loss of ZBP1 expression (Fig. S1C,G). As a control, siRNA-mediated depletion of MLKL prevented both ZBP1- and TNFR1-induced necroptosis (Fig. S1C,D). Together, these data show that endogenous ZBP1 can induce necroptosis in human HT-29 cells.

### The Zα domains and RHIMs of human ZBP1 cooperate to induce necroptosis

To study the contributions of the Zα domains and the RHIMs to ZBP1-mediated necroptosis we transduced HT-29 cells with variants of human ZBP1 using doxycycline-inducible lentivectors (Fig. 1A). These included variants in which both Zα1 and Zα2 domains were mutated (N46A/Y50A and N141A/Y145A) preventing binding to Z-nucleic acids (mutZα1α2), three mutants in which the RHIMs were mutated individually: ^206^IQIG>AAAA (mutRHIM-A), ^264^VQLG>AAAA (mutRHIM-B) or ^332^ATIG>AAAA (mutRHIM-C), proteins only containing the Zα domains (Zα1α2-only) or the RHIMs (RHIMs-only), and the natural splice variant lacking the first Zα domain [ΔZα1 (iso 2)]. Leaky expression from these lentivectors was sufficient to induce ZBP1-dependent cell death upon HSV-1 ICP6^mutRHIM^ infection and (Fig. S1H,I). As controls, the parental HSV-1 strain expressing wild type ICP6 (HSV-1 ICP6^WT^) did not induce detectable levels of cell death and cells stimulated with TNF/BV6/zVAD to induce TNFR1-dependent necroptosis died in a ZBP1-independent manner (Fig. S1I). To be able to monitor ZBP1 localisation upon activation (see below) we fused the ZBP1 variants C-terminally to eGFP-V5, which did not influence its ability to induce cell death compared to proteins only containing short FLAG or V5 tags (Fig. S1H,I).

While wild type ZBP1 induced cell death starting at 8 hours post-infection with HSV-1 ICP6^mutRHIM^, all ZBP1 variants exhibited reduced activities, albeit to varying degrees (Fig. 1B,C). Mutation of both Zα domains (mutZα1α2), which prevents binding to Z-DNA or Z-RNA, or of the first RHIM (mutRHIM-A), which was previously shown to be non-redundant for the recruitment of RIPK1 and RIPK3 to mouse and human ZBP1 (Kaiser *et al*., 2008; Rebsamen *et al*., 2009; Upton *et al*., 2012), completely prevented ZBP1-induced cell death. Deletion of the first Zα domain [ΔZα1 (iso 2)] or mutation of RHIM-B or RHIM-C resulted in intermediate phenotypes displaying on average 40-50 % loss of ZBP1 activity as measured by its capacity to induce cell death (Fig. 1B,C). Truncated ZBP1 proteins only containing the N-terminal Zα domains (Zα1α2-only) or the RHIMs (RHIMs-only) including the C-terminal tail did not exert any activity (Fig. 1B,C). While the previously mentioned ZBP1 domain mutations reduced or prevented cell death following HSV-1 ICP6^mutRHIM^ infection, TNFR1-induced necroptosis remained unaffected (Fig. S1J,K).

Co-immunoprecipitation experiments showed that wild type ZBP1 associated with both RIPK1 and RIPK3 after HSV-1 ICP6^mutRHIM^ infection and with their kinase active forms as shown by autophosphorylation of RIPK1 on Ser166 and RIPK3 on Ser227 (Degterev *et al*, 2008; Sun *et al*., 2012) (Fig. 1D). RIPK3 phosphorylates and activates the pseudokinase MLKL on Thr357 and Ser358 to induce necroptosis (Sun *et al*., 2012). Western blotting showed that wild type human ZBP1 induced phosphorylation of MLKL on Ser358 (Fig. 1D), indicating that ZBP1 activation resulted in the assembly of a necroptotic signalling complex containing activated RIPK1 and RIPK3. Mutation of both Zα domains or of RHIM-A completely prevented the interaction of ZBP1 with RIPK1 and RIPK3 and subsequent MLKL activation, while mutation of RHIM-B or RHIM-C led to reduced but detectable RIPK1/3 recruitment (Fig. 1D). This is in line with the observation that Zα domain or RHIM-A mutation completely prevents HSV-1 ICP6^mutRHIM^-induced cell death, while RHIM-B or RHIM-C mutants still retain some activity (see Fig. 1B,C). The capacity of ZBP1 to induce necroptosis inversely correlated with HSV-1 ICP6^mutRHIM^ replication as measured by the expression of herpesviral *gD*, *ICP27* and *ICP8* transcripts over a 3 day infection period, which were up to 300-fold higher in cells expressing Zα domain or RHIM-A mutant ZBP1 compared to those expressing wild type ZBP1 (Figs. 1E and S1L). Mutations in RHIM-B, RHIM-C or deletion of Zα1 resulted in intermediate phenotypes in line with the partially reduced ability of these ZBP1 variants to induce necroptosis (Figs. 1E and S1L).

Together, these data show that the Zα domains and RHIMs of ZBP1 are required to induce host cell necroptosis and restrict viral replication. Both Zα domains and all three RHIMs contribute to the formation of a necroptotic signalling complex containing ZBP1 and kinase active RIPK1 and RIPK3.

### Zα domain-dependent ZBP1 condensate formation precedes necroptosis

To better understand the kinetic processes involved in ZBP1 activation, we generated HT-29 clones expressing either wild type (clone B9) or Zα domains-mutant (mutZα1α2; clone E6) human ZBP1 fused with a C-terminal eGFP-V5 tag. Both clones showed comparable levels of leaky ZBP1-eGFP expression from the doxycycline-inducible lentivector, which could be further enhanced by doxycycline treatment (Fig. S2A). Similar to their parental polyclonal lines (see Fig. S1I), leaky expression of wild type but not Zα domains-mutant ZBP1 was sufficient to induce cell death after HSV-1 ICP6^mutRHIM^ infection, while both clones responded similarly to TNFR1-induced necroptosis (Fig. S2B).

While human ZBP1 is normally diffusely distributed throughout the cytosol, HSV-1 ICP6^mutRHIM^ infection caused the reorganisation of ZBP1 into foci as early as 5-6 hours post-infection as shown by (live cell) confocal microscopy and imaging flow cytometry (Figs. 2A, C and S2D, movie 1). Hereafter, we refer to these foci as ZBP1 ‘condensates’ describing -in its broadest definition-the local concentration of proteins and nucleic acids into structures that adopt liquid, gel-like or solid states and which are not surrounded by a membrane (Banani *et al*, 2017; Lyon *et al*, 2021). ZBP1 condensate formation preceded the induction of cell death, which commenced 8 hours post-infection (Figs. 2C, S2B, movie 1). Both the number and size and of these ZBP1 condensates increased over time (Figs. 2B and S2D) and their formation depended on the presence of functional Zα domains (Figs. 2D,E and S2E, movie 2). The induction of ZBP1 condensates did not depend on the induction of necroptosis as infection with an HSV-1 strain expressing wild type ICP6, which blocks necroptosis (see Fig. S1I, movie 3), also resulted in the formation of ZBP1 foci that were indistinguishable in both number and size compared to those formed after HSV-1 ICP6^mutRHIM^ infection (Figs. 2D,E and S2C,E). Also infection with influenza A virus (IAV), which induces both ZBP1-dependent apoptosis and necroptosis in mouse and human cells (Kuriakose *et al*., 2016; Thapa *et al*., 2016; Zhang *et al*., 2020), resulted in Zα domains-dependent ZBP1 condensation and cell death (Fig. S2F,G). Only cells that were productively infected with IAV displayed ZBP1 foci suggesting that ZBP1 condensate formation is a cell-intrinsic process (Fig. S2G).

**Figure 2.**
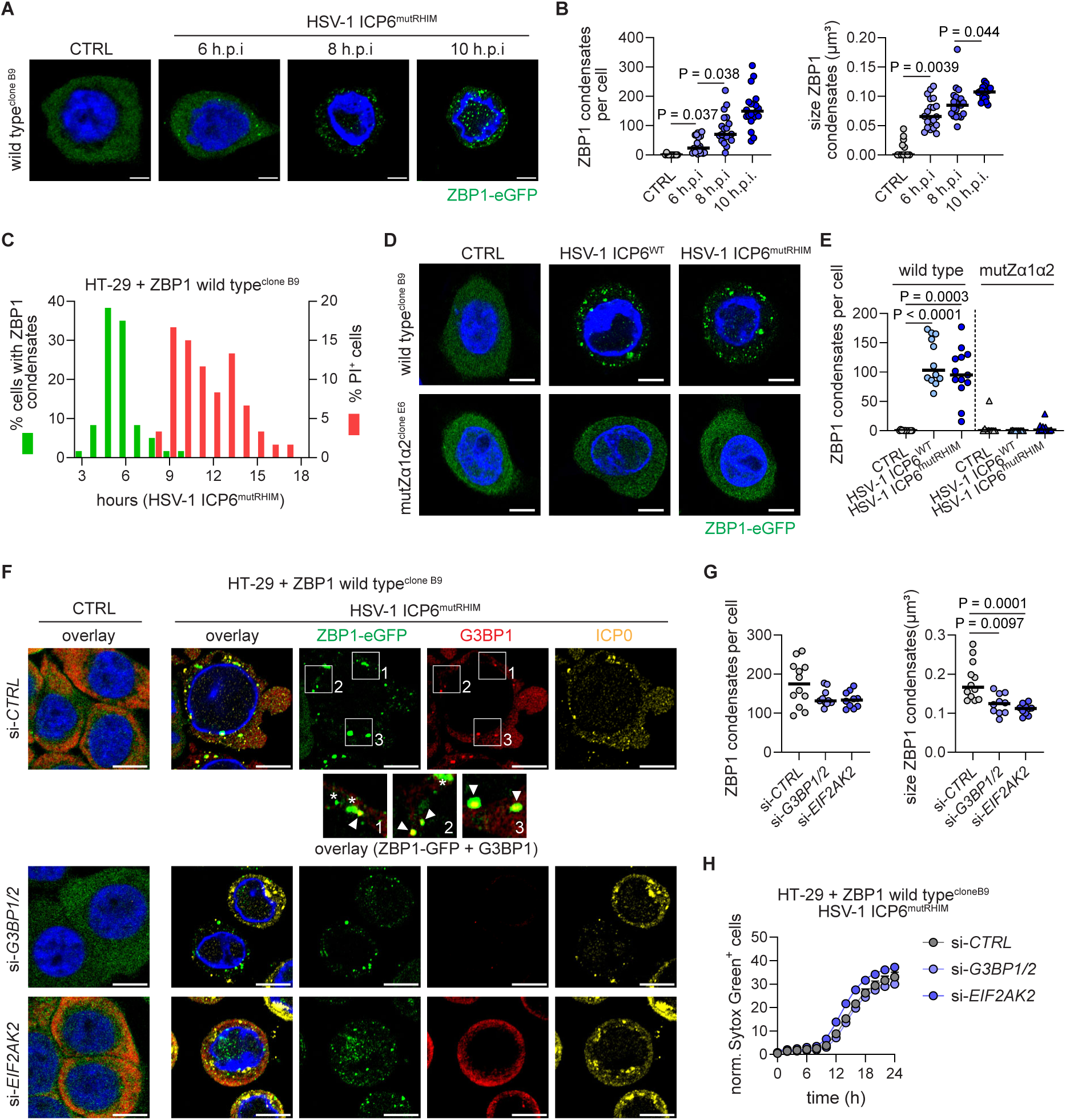
ZBP1 forms condensates and necroptosis induction does not depend on stress granules. **(A,B)** HT-29 cells expressing wild type (isoform 1) human ZBP1-eGFP-V5 (clone B9) were infected with HSV-1 ICP6^mutRHIM^ (MOI of 5). Cells were analysed by confocal microscopy at different hours post-infection (h.p.i.). **(A)** Representative images of ZBP1 visualised using its eGFP tag (green) and DAPI (blue). Scale bars, 5 µm. **(B)** Quantification of the number of ZBP1-eGFP condensates per cell (left graph) and condensate size (right graph). ZBP1 condensates were analysed in 3D-images and every dot represents a z-stack. P values by One-Way ANOVA. **(C)** HT-29 cells expressing wild type (isoform 1) human ZBP1-eGFP-V5 (clone B9) were infected with HSV-1 ICP6^mutRHIM^ (MOI of 5). The percentage of cells (n = 60) containing ZBP1-eGFP condensates and PI uptake were measured by live cell imaging. **(D,E)** HT-29 cells expressing wild type (isoform 1) human ZBP1-eGFP-V5 (clone B9) or mutZα1Zα2 human ZBP1-eGFP-V5 (clone E6) were infected with HSV-1 ICP6^mutRHIM^ or HSV-1 ICP6^WT^ (MOI of 5) for 9 hours. **(D)** Representative images of ZBP1 visualised using its eGFP tag (green) and DAPI (blue). Scale bars, 5 µm. **(E)** Quantification of ZBP1-eGFP-V5-positive condensates per cell. P values by One-Way ANOVA. ZBP1 condensates were analysed in 3D-images and every dot represents a z-stack. **(F-H)** HT-29 cells expressing wild type (isoform 1) human ZBP1-eGFP-V5 (clone B9) or mutZα1Zα2 human ZBP1-eGFP-V5 (clone E6) were transfected with siRNAs targeting *G3BP1* and *G3BP2*, or *EIF2AK2* or non-targeting control siRNAs (si-*CTRL*). 48 hours later, cells were infected with HSV-1 ICP6^mutRHIM^ (MOI of 5) for 9 hours. **(F)** Representative images showing ZBP1 (green), G3BP1 (red), DAPI (blue) and the HSV-1 protein ICP0 (yellow). The three zoomed areas depict overlays of ZBP1-eGFP and G3BP1. Scale bars, 10 µm. **(G)** Quantification of the number of ZBP1-eGFP condensates per cell (left graph) and measurement of condensate size (right graph). ZBP1 condensates were analysed in 3D-images and every dot represents a z-stack. P values by One-Way ANOVA. **(H)** Cell death was quantified by measuring Sytox green uptake every 2 hours using Incucyte cell imaging. The number of Sytox green^+^ cells per image at each time point was divided by the percentage of confluency to obtain normalised values plotted as “norm. Sytox geen^+^ cells” on the Y-axis. Lines represent a sigmoidal, 4PL fit.

Arsenite-induced oxidative stress induces Zα domains-dependent translocation of ZBP1 to stress granules (Deigendesch *et al*., 2006; Ng *et al*, 2013), which can serve as activating sites to induce necroptosis (Szczerba *et al*, 2023; Yang *et al*, 2023). HSV-1 infection also induces stress granule formation, a process that depends on translational inhibition by the dsRNA sensor Protein kinase R (PKR) through eIF2α phosphorylation (Dauber *et al*, 2016). We therefore tested whether ZBP1 condensates colocalised with stress granules. Although most G3BP1-positive stress granules that formed after HSV-1 ICP6^mutRHIM^ infection colocalized with ZBP1 foci (Fig. 2F, white arrows), not all ZBP1 condensates localized with stress granules (Fig. 2F, white *). siRNA-mediated knockdown of either the essential stress granule forming proteins G3BP1 and G3BP2 or of the stress granule-inducing dsRNA sensor PKR (encoded by *EIF2AK2*) did not reduce the number of ZBP1 condensates per cell although their size was reduced (Figs. 2F,G and S2H,I). Moreover, knockdown of G3BP1/2 or PKR did not prevent ZBP1-mediated necroptosis neither in settings of ectopic nor of endogenous ZBP1 expression (Figs. 2H and S2J,K).

In sum, HSV-1 infection resulted in the formation of ZBP1 condensates, only some of which colocalized with stress granules. Stress granules formation, however, was not functionally relevant for the induction of virus-induced cell death. The reorganisation of ZBP1 in condensates preceded cell death, suggesting this process is an early event in necroptosis induction.

### Z-RNA-Zα domain interactions induce the formation of ZBP1 condensates

To understand how the Z-nucleic acid-interacting Zα domains contribute to ZBP1 condensate formation we stained HSV-1 ICP6^WT^-infected cells with a monoclonal antibody (clone Z22) that binds to both Z-DNA and Z-RNA (Moller *et al*, 1982; Zhang *et al*., 2020). It should be noted that the Z22 antibody, similar to Zα domains, can also stimulate Z-prone sequences to form Z-DNA (Moller *et al*., 1982). Nevertheless, the presence of Z22-positive nucleic acids serves as a proxy for the presence of possible ZBP1 activating nucleic acids in an infected cell. The Z22 antibody stained the cytosol of HT-29 cells as early as 5 hour after HSV-1 ICP6^WT^ infection and this signal increased over time (Fig. 3A,B). Treatment of HSV-1 ICP6^WT^-infected cells with RNase A, which cleaves ss- and dsRNA, before staining reduced the Z22 signal (Figs. 3C and S3A). Pre-treatment with DNase I, which degrades ss- and dsDNA, had no measurable impact of Z22 signal (Figs. 3C and S3A). This indicates that Z-RNA and/or Z-RNA-prone structures accumulate in HSV-1-infected cells. As controls, RNase A treatment reduced the presence of A-form dsRNA structures accumulating after HSV-1 infection as detected by the dsRNA-specific J2 antibody (Weber *et al*, 2006) and DNase I treatment prevented genomic DNA staining by DAPI (Fig. S3A). As previously shown (Zhang *et al*., 2020), infection with IAV also resulted in an increase in Z22 signal in the cytoplasm (Fig. S3B,C).

**Figure 3.**
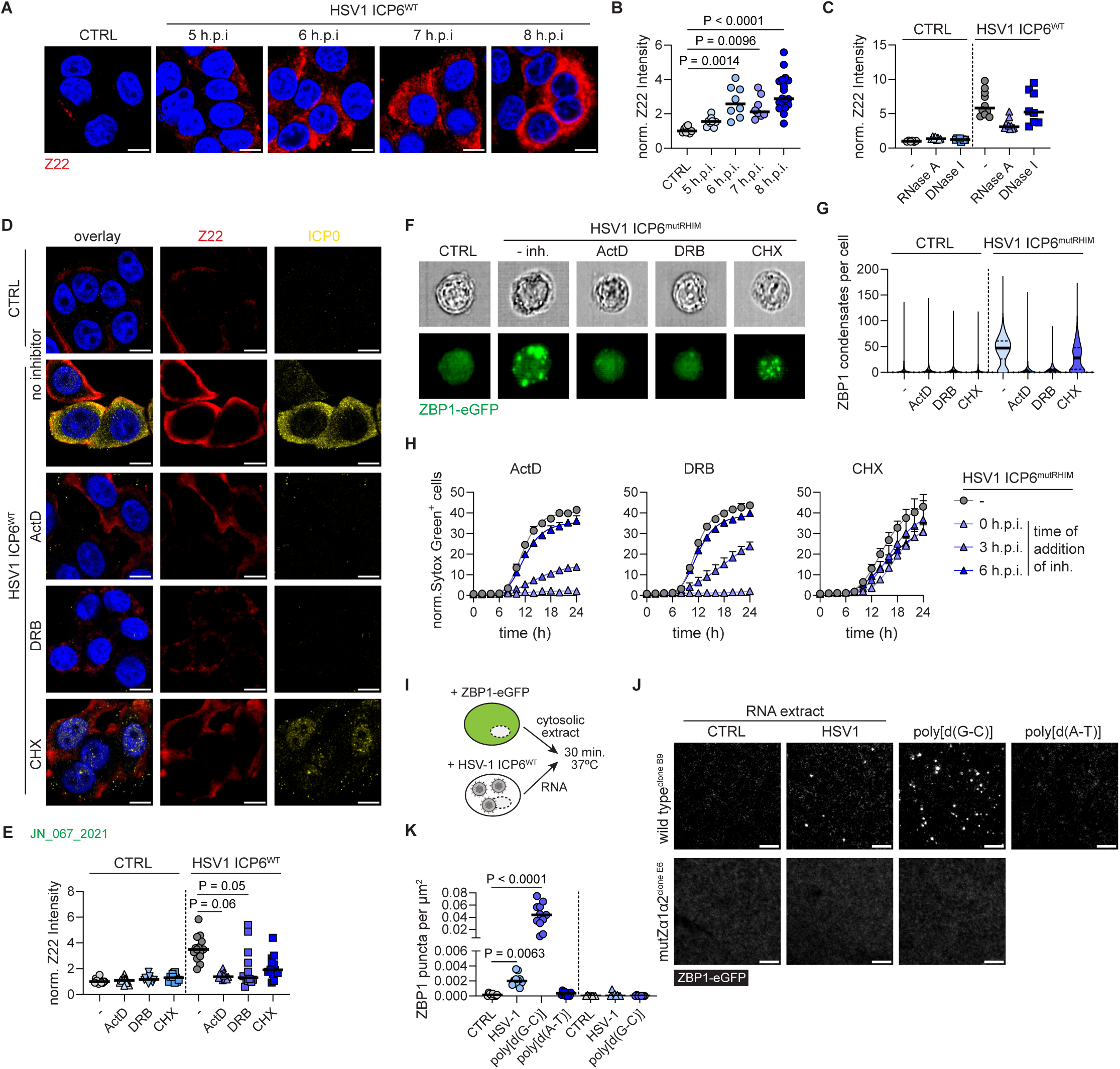
Z-RNA-Zα domain interactions induce ZBP1 condensate formation. **(A,B)** HT-29 cells were not infected (CTRL) or were infected with HSV-1 ICP6^WT^ (MOI of 5). At different hours post-infection (h.p.i.), cells were stained with an antibody recognising Z-prone or Z-RNA/DNA (Z22). **(A)** Representative images showing Z22 (red) and DAPI (blue) staining. **(B)** Quantification of mean fluorescent intensity of Z22, normalised to the Z22 signal in non-infected (CTRL) cells, plotted as “norm. Z22 intensity” on the Y-axis. Each dot represents a single cell. Scale bars, 10 µm. P values by One-Way ANOVA. **(C)** HT-29 cells were infected with HSV-1 ICP6^WT^ (MOI of 5) for 6 hours. Cells were treated with RNase A or DNase I before Z22 staining. The Z22 signal was quantified as in (B). **(D-G)** HT-29 cells were left untreated (no inhibitor) or treated with 5 μM actinomycin D (ActD), 50 μM RNA polymerase II inhibitor 5,6-dichlorobenzimidazole riboside (DRB) or 1 μg/ml cycloheximide (CHX). 30 min. later, cells were then infected for 6 hours with HSV-1 ICP6^WT^ (MOI of 5) (D,E) or 9 hours (F,G) with HSV-1 ICP6^mutRHIM^ (MOI of 5). **(D)** Representative images showing Z22 (red), the HSV-1 protein ICP0 (yellow) and DAPI (blue). Scale bars, 10 µm. **(E)** The Z22 signal was quantified as in (B). P values by One-Way ANOVA. **(F,G)** Cells were analysed using imaging flow cytometry and ZBP1-eGFP condensates were quantified (G). Representative brightfield and ZBP1-eGFP images are shown in (F). **(H)** HT-29 cells expressing wild type (isoform 1) human ZBP1-eGFP-V5 (clone B9) were infected with HSV-1 ICP6^mutRHIM^ (MOI of 5). Cells were left untreated (-) or were treated at 0, 3 or 6 h.p.i. with 5 μM ActD, 50 μM DRB or 1μg/ml CHX. Cell death was quantified by measuring Sytox green uptake every 2 hours. The number of Sytox green^+^ cells per image at each time point was divided by the percentage of confluency to obtain normalised values plotted as “norm. Sytox green^+^ cells” on the Y-axis. Lines represent a sigmoidal, 4PL fit. **(I)** Schematic representation of the *in vitro* complementation assay. Cytosolic extracts from HT-29 expressing human ZBP1-eGFP-V5 are incubated with RNA isolated from HT-29 cells infected with HSV-1 ICP6^WT^ (MOI of 5) and incubated for 30 minutes at 37 °C. ZBP1-eGFP condensates are then imaged by total internal reflection microscopy. **(J,K)** *In vitro* complementation assay combining of cytosolic lysates from HT-29 cells expressing wild type (isoform 1) or mutZα1Zα2 human ZBP1-eGFP-V5 with RNA isolated from uninfected (CTRL) or HSV-1 ICP6^WT^-infected (MOI of 5) cells. As controls, cytosolic extracts were incubated with Z-prone DNA, poly[d(G-C)], or B-prone DNA, poly[d(A-T)]. **(J)** Representative images depicting ZBP1-eGFP puncta (white) and **(K)** Quantification of ZBP1-eGFP-V5 condensates per square μm. Scale bars, 5 µm. P values by One-Way ANOVA.

In the context of infection with murine cytomegalovirus (Maelfait *et al*., 2017; Sridharan *et al*., 2017), HSV-1 (Guo *et al*., 2018) or the vaccinia poxvirus (Koehler *et al*, 2021) host RNA polymerase II-mediated transcriptional activity is required to activate ZBP1, independently of protein translation or viral genomic DNA replication. This indicates that accumulation of immediate early or early viral and/or altered host transcripts promotes the formation of Z-RNA or Z-RNA prone structures that activate ZBP1. Indeed, addition of the RNA polymerase II inhibitor 5,6-dichlorobenzimidazole riboside (DRB) or of the general transcriptional inhibitor actinomycin D (ActD), which both inhibited the expression of the viral immediate early protein ICP0 (Figs. 3D and S3D), reduced the Z22 signal in HSV-1 ICP6^WT^-infected HT-29 cells (Fig. 3D,E). Transcriptional inhibition by DRB or ActD also prevented the formation of ZBP1 condensates and inhibited cell death induction in ZBP1-eGFP expressing HT-29 cells infected with HSV-1 ICP6^mutRHIM^ (Fig. 3F-H). Inhibition of RNA polymerase activity remained partially effective in preventing cell death at 3 hours post-infection and had lost its effect when DRB or ActD was added at 6 hours post-infection (Fig. 3H). In contrast, blockade of translation by cycloheximide (CHX), which also reduced ICP0 protein production (Figs. 3D and S3D), did not significantly impact on Z22 antibody staining, ZBP1 condensate formation or necroptosis induction (Fig. 3D-H).

Z-RNA or Z-RNA-prone nucleic acids accumulate as early as 5 hours post-infection and this coincided with the appearance of ZBP1 foci, suggesting that the interaction of Z-nucleic acids with ZBP1 stimulated condensate formation. To test this hypothesis we developed an *in vitro* complementation assay in which we incubated a ZBP1-eGFP-containing cytosolic extract from non-infected cells with a Z-prone dsDNA polymer, poly[d(G-C)], consisting out of alternating guanosine and cytidine nucleotides for 30 minutes at 37°C and monitored ZBP1 condensate formation using total internal reflection fluorescence microscopy (Fig. 3I). Incubation of ZBP1-eGFP-containing lysates with poly[d(G-C)], but not with poly[d(A-T)], a B-DNA polymer consisting of alternating adenosines and thymidines, caused ZBP1 puncta formation in the *in vitro* assay and this required functional Zα domains (Fig. 3J-K). Similarly, transfection of the Z-prone sequence poly[d(G-C)] but not B-form poly[d(A-T)] resulted in the formation of ZBP1 foci in HT-29 cells (Fig. S3E,F). *In vitro* incubation of ZBP1-eGFP lysates with RNA extracted from HSV-1 ICP6^WT^-infected HT-29 cells, but not from non-infected cells, caused ZBP1 condensate formation (Fig. 3I-K) showing that HSV-1 infection resulted in the accumulation of ZBP1-interacting RNA sequences. Together, these data show that the interaction of ZBP1’s Zα domains with Z-RNA or Z-prone RNA molecules that accumulate during HSV-1 infection causes ZBP1 condensate formation.

### ZBP1 condensates form independently of the RHIMs and RIPK1/3

RHIMs, including those from ZBP1, RIPK1, RIPK3 and TRIF, have the capacity to polymerise into β-amyloidal fibrils (Baker *et al*, 2020). We therefore asked whether the RHIMs of ZBP1 and/or those of RIPK1/3 stimulated condensate formation. Confocal microscopy and imaging flow cytometry, however, showed that individual mutation of neither RHIM-A, RHIM-B nor RHIM-C affected ZBP1-eGFP condensate formation upon infection with HSV-1 expressing either wild type or RHIM-mutant ICP6 (Figs. 4A,B and S4A,B). A ZBP1 protein only containing the N-terminal Zα domains (Zα1α2-only) was still able to form a similar amount of foci per cell showing that the formation of ZBP1 condensates can occur independently of the RHIMs. In contrast, mutation (mutZα1α2) or removal (RHIMs-only) of both Zα domains prevented HSV-1-induced ZBP1 condensate formation after HSV-1 ICP6^WT^ or HSV-1 ICP6^mutRHIM^ infection (Figs. 4A,B and S4A,B). These results were confirmed by the *in vitro* complementation assay: incubation of cytosolic extracts of cells expressing RHIM-A, -B or -C-mutant ZBP1-eGFP with RNA from HSV-1-infected cells promoted the formation of ZBP1-eGFP condensates (Fig. 4C,D). Cytosolic extracts from cells expressing a ZBP1-eGFP variant containing only the Zα domains (Zα1α2-only) still supported ZBP1 condensate formation albeit less efficiently (Fig. 4C,D). Neither siRNA-mediated depletion of RIPK1 or RIPK3 expression nor inhibition of their kinase activities affected ZBP1 condensate formation in HT-29 cells after infection with either wild type of ICP6 RHIM-mutant HSV-1 (Fig. S4C-E). Together, these data show that neither the RHIMs of ZBP1 nor the RHIM-containing downstream signalling proteins RIPK1 or RIPK3 are essential to mediate the formation of ZBP1 condensates after HSV-1 infection.

**Figure 4.**
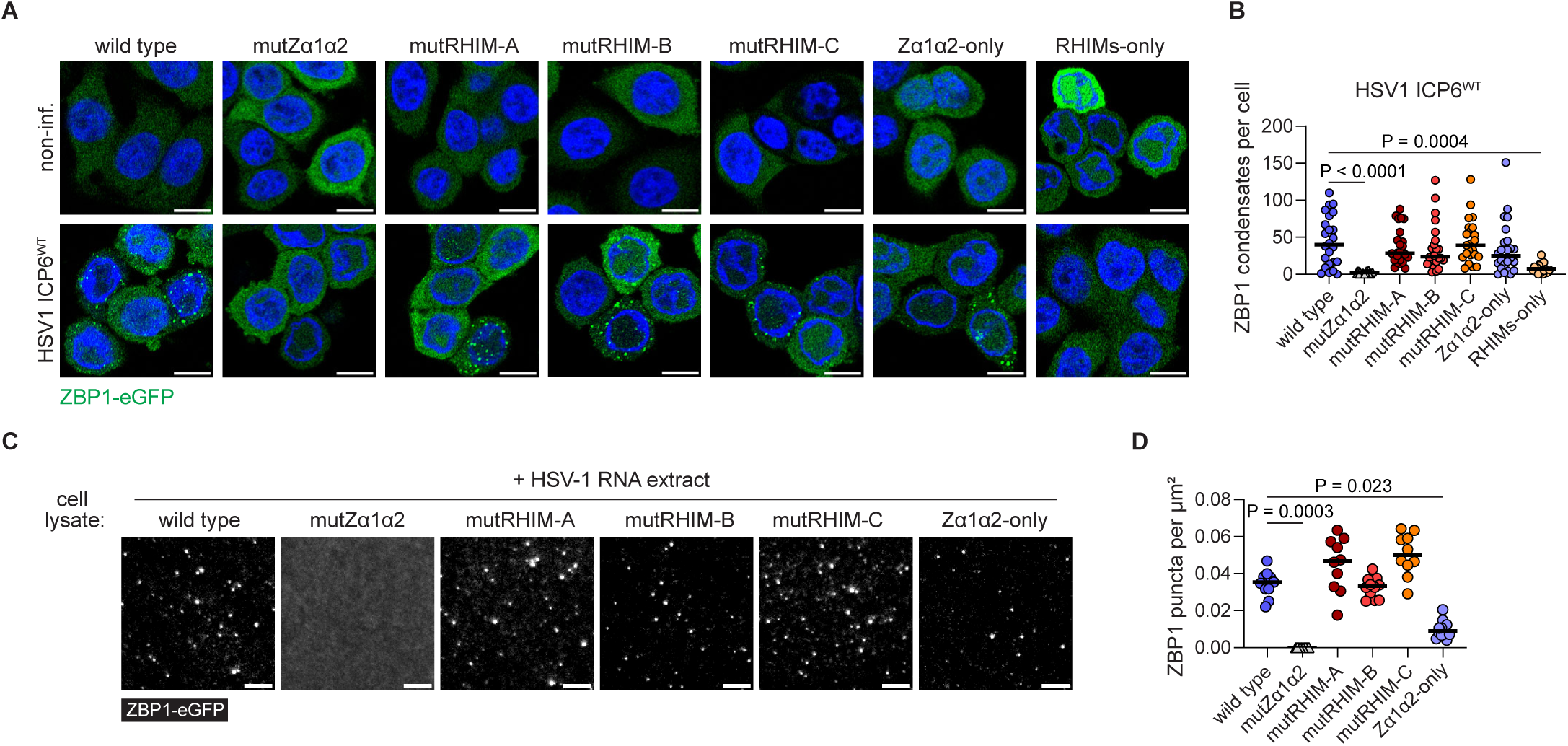
ZBP1 condensates form independently of the RHIMs. **(A,B)** HT-29 cells transduced with doxycycline-inducible lentivectors expressing the indicated human ZBP1-eGFP-V5 variants were infected with HSV-1 ICP6^WT^ at a multiplicity of infection (MOI) of 5. 9 hours post-infection cells were analysed by confocal microscopy. **(A)** Representative images showing ZBP1-eGFP (green) and DAPI (blue). Scale bars, 10 µm. **(B)** Quantification of the number of ZBP1-eGFP condensates per cell. ZBP1 condensates were analysed in 3D-images and every dot represents a single cell. P value by One-Way ANOVA. **(C,D)** *In vitro* complementation assay combining cytosolic lysates from HT-29 cells expressing the indicated human ZBP1-eGFP-V5 variants with RNA isolated from HT-29 cells infected with HSV-1 ICP6^WT^ (MOI of 5). **(C)** Representative images depicting ZBP1-eGFP puncta (white). **(D)** Quantification of ZBP1-eGFP-V5 puncta per square μm. Scale bars, 5 µm. P values by One-Way ANOVA.

### The RHIMs of ZBP1 promote the assembly of solid state condensates

To analyse the dynamics and material state of the ZBP1 condensates we next used live cell confocal microscopy and fluorescent recovery after photobleaching (FRAP). Tracking of ZBP1 condensates that formed in cells after infection with HSV-1 showed that larger condensates (> 0.17 µm^2^) were less mobile than smaller foci, suggesting that ZBP1 condensates progressively mature into larger less-dynamic structures (Fig. S5A). In the large immobile foci, we did not observe any recovery of ZBP1-eGFP fluorescence indicating a solid state of these condensates with no molecules moving ‘in or out’ of these structures (Figs. 5A,B and S5B,C, movie 4). The RHIM of ICP6 did not interfere with the dynamics or state of these ZBP1 assemblies since infection with an HSV-1 strain expressing RHIM-mutant ICP6 did not change the speed or the FRAP profile of ZBP1 condensates compared to infection with a wild type virus (Fig. S5A-C, movie 5). We then monitored the contribution of the Zα domains and of the RHIMs to the biophysical characteristics of ZBP1 condensates. Interestingly, mutation of RHIM-A (mutRHIM-A) resulted in partial recovery of the ZBP1-eGFP fluorescent signal, while mutation of RHIM-B or -C (mutRHIM-B or -C) or deletion of the first Zα domain [ΔZα1 (iso 2)] did not change the material state of ZBP1 condensates (Fig. 5A,B, movies 6-9 and 11). Moreover, ZBP1 assemblies formed by a protein variant containing only the Zα domains and lacking all RHIMs (Zα1α2-only) displayed even better recovery after photobleaching (Fig. 5A,B, movie 10). Together, these data show that the Zα domains promote the formation of ZBP1 condensates that remain partially dynamic, while the RHIMs are required to assemble solid state structures, compatible with their capacity to form amyloids.

**Figure 5.**
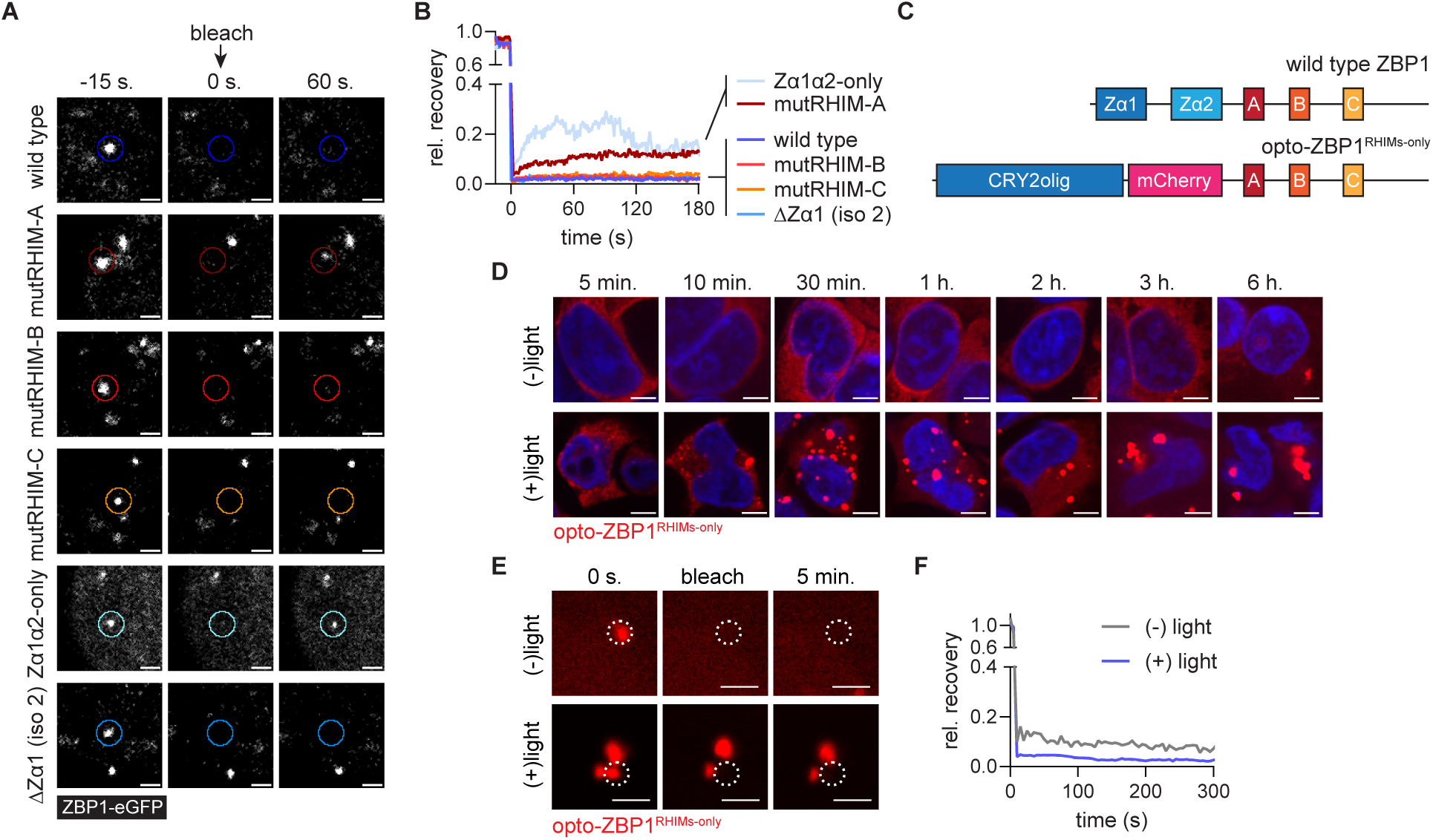
The RHIMs of ZBP1 support the assembly of solid state condensates. **(A,B)** HT-29 cells transduced with doxycycline-inducible lentivectors expressing the indicated human ZBP1-eGFP-V5 variants were infected with HSV-1 ICP6^WT^ at a multiplicity of infection (MOI) of 5. **(A)** Representative images of ZBP1 condensates, before (-15 s.), immediately after (0 s.) or 60 seconds (60 s.) after photobleaching. The bleached areas are highlighted with a coloured circle. Scale bars, 2 µm. **(B)** Fluorescent recovery after photobleaching (FRAP) of ZBP1 condensates (n ≥ 4) formed by the indicated human ZBP1 variants. The fluorescent intensity of the photobleached area at the indicated time point was normalised to the average fluorescent intensity at -15 s., which was set at 1 and plotted at as “rel. recovery” in the Y-axis. **(C)** Schematic overview of wild type (isoform 1) and the optogenetic ZBP1 protein (opto-ZBP1^RHIMs-only^). In opto-ZBP1^RHIMs-only^ the Zα domains were replaced by a self-oligomerising CRY2olig domain, enabling Zα domain-independent ZBP1 clustering, and an mCherry fluorescent tag allowing protein visualisation. The protein also contains a C-terminal His- and FLAG-tag (not shown). **(D-F)** Flp-In 293 T-REx cells expressing opto-ZBP1^RHIMs-only^ under a doxycycline-inducible promotor were treated with 1 µg/ml doxycycline for 24 hours. Cells were then kept in the dark [(-)light)] or exposed to 10 V blue light [(+)light)] for 2 min. **(D)** After the indicated time points, cells were fixed and analysed by confocal microscopy. Representative images showing opto-ZBP1^RHIMs-only^ (red) and DAPI (blue). Scale bars, 5 µm. **(E)** Representative images of opto-ZBP1^RHIMs-only^ foci, before (0 s.), immediately after (bleach) or 5 minutes (5 min.) after photobleaching. The bleached areas are highlighted with a dotted circle. Scale bars, 2 µm. **(F)** FRAP curves of opto-ZBP1 foci (n ≥ 6) that formed either spontaneously [(-) light)] or 3 hours after blue light exposure [(+) light)]. The fluorescent intensity of the photobleached area at the indicated time point was normalised to the average fluorescent intensity at 0 s., which was set at 1, and plotted at as “rel. recovery” in the Y-axis.

To further assess the potential of the RHIMs of ZBP1 to form stable solid state condensates without interference from the Zα domains, we generated Flp-In 293 T-REx cells expressing an optogenetic ZBP1 construct (opto-ZBP1^RHIMs-only^) in which we replaced the Zα domains with a CRY2olig domain coupled to mCherry under control of a doxycycline-inducible promotor (Figs. 5C, S5D). The CRY2olig domain normally forms reversible homo-oligomeric complexes upon exposure to blue light (Taslimi *et al*., 2014). We employed this system to promote homotypic interactions between the RHIMs of individual ZBP1 molecules in a ligand-independent manner. Doxycycline treatment resulted in a diffuse cytosolic expression pattern of opto-ZBP1^RHIMs-only^ and exposure to blue light induced its redistribution into abundant cytosolic clusters within 10 minutes after light-induced oligomerisation (Fig. S5E). In cells exposed to blue light (10 V, 2 min.) we observed a slow and progressive assembly of opto-ZBP1 into cytoplasmic foci (Fig. 5D, movie 12). The first sizeable structures became visible approximately 10 min. after the blue light pulse. At later time points the foci coalesced into larger less mobile aggregates that remained very stable (Fig. 5D, movie 12). Spontaneous formation of opto-ZBP1^RHIMs-only^ foci was occasionally observed in cells that were not exposed to blue light (Fig. 5D). This is probably mediated through spontaneous RHIM-RHIM interactions resulting from opto-ZBP1^RHIMs-only^ overexpression. Indeed, doxycycline-induced overexpression of a ZBP1 variant only containing the RHIMs (RHIMs-only) in HT-29 cells similarly resulted in the accumulation of large complexes, bypassing the need to need for initial ZBP1 concentration mediated by the interaction between Z-nucleic acids and the Zα domains (see Fig. S4F). Finally, we analysed the material state of the large opto-ZBP1^RHIMs-only^ condensates using FRAP. Both opto-ZBP1^RHIMs-only^ foci that formed spontaneously and those that formed upon blue light exposure did not recover upon photobleaching, indicating a solid state (Fig. 5E,F, movies 13 and 14) similar to the ZBP1-eGFP condensates that formed after HSV-1 infection (see Figs. 5A,B and S5B,C). Together, these data show that the RHIMs of ZBP1 promote the stabilisation of ZBP1 condensates into solid state structures.

### ZBP1 forms amyloidal signalling complexes

β-amyloidal fibrils, including those formed by the RHIMs of RIPK1 and RIPK3, are resistant to high concentrations of denaturing agents such as sodium dodecyl sulfate (SDS) or urea (Baker *et al*., 2020; Li *et al*., 2012; Mompean *et al*., 2018). To determine the possible amyloidal nature of the ZBP1 condensates we performed semi-denaturing detergent agarose gel electrophoresis (SDD-AGE), which assesses the stability of protein complexes in the presence of 2 % SDS (Liu *et al*, 2017). This showed that ZBP1 assembled into SDS-resistant oligomers following infection with ICP6 RHIM-mutant HSV-1 (Fig. 6A). These ZBP1 complexes were also resistant to 8 M Urea, but not to heat denaturation, further supporting their amyloidal nature (Fig. 6B). As a control, ZBP1 did not oligomerise after the induction of TNFR1-induced necroptosis (Figs. 6A). TNFR1-induced activation of necroptosis depends on the assembly of an amyloidal RIPK1/3 signalling platform that activates MLKL (Chen *et al*., 2022; Li *et al*., 2012; Liu *et al*., 2017). Similarly, both RIPK1 and RIPK3 formed SDS-resistant oligomers after ZBP1-mediated necroptosis induced by infection with HSV-1 ICP6^mutRHIM^ (Fig. 6A).

**Figure 6.**
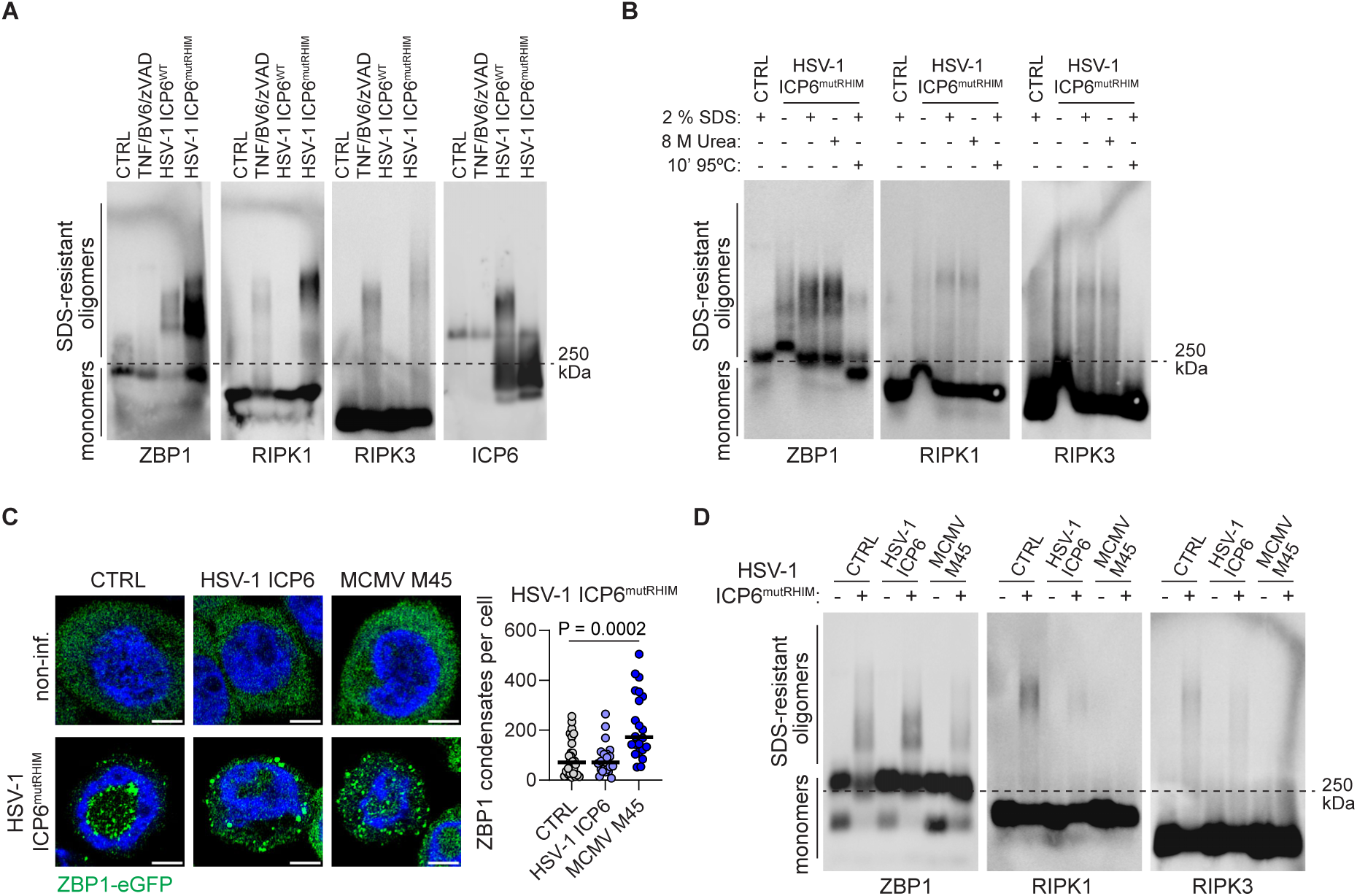
ZBP1 forms an amyloidal signalling complex and ICP6 inhibits downstream RIPK1/3 oligomerisation. **(A,B)** HT-29 cells expressing wild type (isoform 1) human ZBP1-eGFP-V5 (clone B9) were left untreated (CTRL), infected with HSV-1 ICP6^WT^ or HSV-1 ICP6^mutICP6^ (MOI of 5) for 9 hours or stimulated with 30 ng/ml TNF, 20 μM zVAD and 5 μM BV6 for 4 hours. **(A)** Cell lysates were analysed by semi-denaturing detergent agarose gel electrophoresis (SDD-AGE). **(B)** Cell lysates were treated with either with 2 % SDS, 8 M urea or heat (10 min. at 95°C) and analysed by SDD-AGE. **(C,D)** HT-29 cells expressing wild type (isoform 1) human ZBP1-eGFP-V5 (clone B9) were transduced with lentivectors expressing the viral RHIM-containing proteins HSV-1 ICP6 or MCMV M45 and the cells were infected with HSV-1 ICP6^mutRHIM^ (MOI of 5) for 9 hours. **(C)** Left: representative images show ZBP1 condensates (green) and DAPI (blue). Scale bars, 10 μm. Right: Quantification of the number of ZBP1-eGFP condensates per cell. ZBP1 condensates were analysed in 3D-images and every dot represents a z-stack. P value by One-Way ANOVA. **(D)** Cell lysates were analysed by SDD-AGE. The dotted line in (A), (B) and (C) indicates the 250 kDa molecular weight marker.

Oligomerisation of ZBP1, RIPK1 and RIPK3 after HSV-1 ICP6^mutRHIM^ infection was blocked by inhibition of transcription by ActD or DRB, but less so by inhibition of protein synthesis by CHX (Fig. S6A). This is in line with the observation that the formation of ZBP1 condensates is dependent on the interaction of newly synthesized RNA with the Zα domains of ZBP1 (see Fig. 3F,G). Indeed, Zα domains-mutant ZBP1 (mutZα1α2) does not form SDS-resistant higher-order structures (see Fig. S7F). Together, these data show that while the Zα domains of ZBP1 initiate the formation of ZBP1 condensates, the RHIMs are required to establish solid state and SDS-resistant amyloidal ZBP1 oligomers, that support the assembly of an amyloidal ZBP1-RIPK1-RIPK3 signalling complex.

### ICP6 inhibits ZBP1-dependent RIPK1/3 oligomerisation

We previously showed that ICP6 does not influence the kinetics of ZBP1 condensate formation (see Figs. 2D,E and 2E) nor their physical state (see Fig. S5A-C), suggesting that ICP6 acted downstream of ZBP1 oligomerisation. Indeed, while infection with HSV-1 expressing wild type ICP6 completely prevented RIPK1 and RIPK3 oligomerisation and blocked the induction of necroptosis as indicated by the absence of RIPK1 Ser166, RIPK3 Ser227 and MLKL Ser358 phosphorylation (Fig. S6B), it only partially inhibited the formation of SDS-resistant ZBP1 oligomers (Figs. 6A and S7F). Similar to the host RHIM-containing proteins ZBP1, RIPK1 and RIPK3, ICP6 also formed SDS-resistant oligomers and this depended on its RHIM (Fig. 6A). To test whether other viral RHIM-containing proteins interfered with ZBP1-induced necroptosis signalling in a similar manner, we expressed M45, a RHIM-containing homologue of ICP6 encoded by murine cytomegalovirus (MCMV), in ZBP1-eGFP expressing HT-29 cells. As a control we also generated cells that expressed ICP6. While ectopic expression of both M45 and ICP6 efficiently inhibited ZBP1 and TNFR1-induced necroptosis (Fig. S6C,D), they did not interfere with the formation of ZBP1 condensates (Fig. 6C) and they did not or only partially prevented the assembly of SDS-resistant ZBP1 oligomers (Fig. 6D). Instead, M45 and ICP6 blocked RIPK1 and RIPK3 oligomerisation (Fig. 6D), consistent with the our observation with virally encoded ICP6 (see Figs. 6A and S7F) and suggesting that these viral RHIM-containing proteins predominantly inhibit ZBP1 signalling downstream of stable ZBP1 oligomerisation.

### RIPK1 kinase activity induces ZBP1-dependent RIPK1/RIPK3 oligomerisation

In mouse cells virus-induced ZBP1-mediated necroptosis can proceed independently of RIPK1 (Nogusa *et al*, 2016; Upton *et al*., 2012), while in human cells this requires RIPK1’s enzymatic activity (Amusan *et al*., 2025). Indeed, CRISPR-Cas9-mediated deletion of RIPK1 or inhibition of its enzymatic activity by Nec-1s confirmed that the induction of necroptotic cell death by ZBP1 depended on RIPK1 and its kinase activity in human cells, but not in mouse cells (Figs. 7A,B and S2A-C). As controls, both RIPK1 depletion and Nec-1s treatment prevented TNFR1-induced necroptosis in both human and mouse cells (Fig. S7D,E). As opposed to Nec-1s treatment, genetic removal of RIPK1 did not fully prevent HSV-1 ICP6^mutRHIM^-mediated cell death in human HT-29 cells, particularly at later time points after infection (Figs. 7A and S7B). The remaining cell death was blocked by addition of the RIPK3 inhibitor GSK’840, but not Nec-1s (Figs. 7A and S7B), suggesting that in conditions of RIPK1 deficiency, human ZBP1 can assemble a necroptotic signalling complex independently of RIPK1 albeit less efficiently.

**Figure 7.**
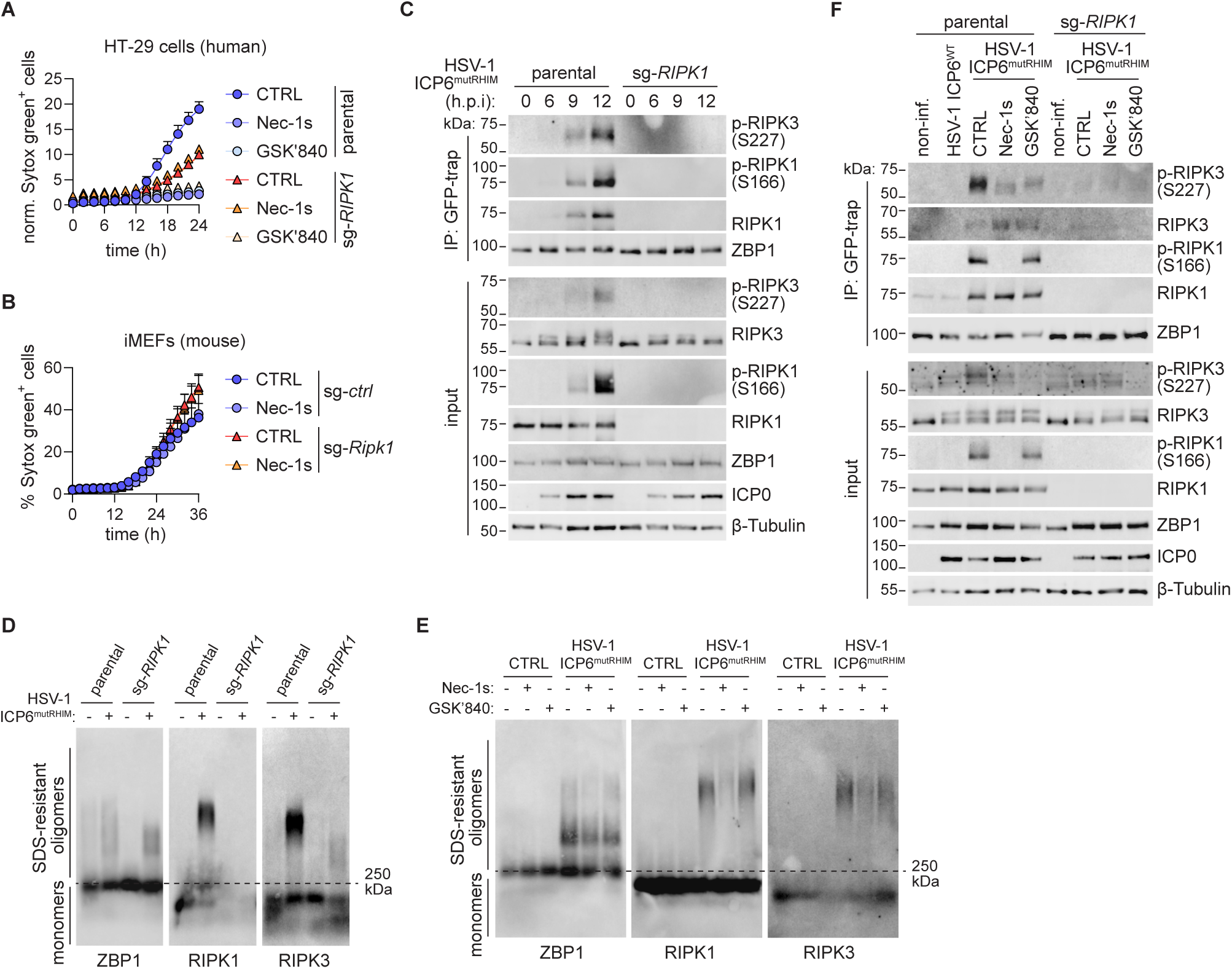
The kinase activity of RIPK1 is required for RIPK1/3 oligomerisation downstream of human ZBP1. **(A)** Parental (clone B9) and sg-*RIPK1* HT-29 cells expressing the eGFP-V5-tagged human ZBP1 were pre-treated with the RIPK1 kinase inhibitor Nec-1s (5 µM) or RIPK3 kinase inhibitor GSK’840 (3 µM) for 30 minutes and then infected with HSV-1 ICP6^mutRHIM^ (MOI of 5). Cell death was quantified by measuring Sytox green uptake every 2 hours using Incucyte cell imaging. The number of Sytox green^+^ cells per image at each time point was divided by the percentage of confluency to obtain normalised values plotted as “norm. Sytox green^+^ cells” on the Y-axis. Lines represent a sigmoidal, 4PL fit. **(B)** Sg-*ctrl* and sg-*Ripk1* immortalised mouse fibroblasts (iMEFs) were pre-treated with Nec-1s (5 µM) for 30 minutes and then infected with HSV-1 ICP6^mutRHIM^ (MOI of 5). Cell death was quantified by measuring Sytox green uptake. The number of Sytox green^+^ cells per image at each time point was divided by the number of Sytox green^+^ cells per image of triton X-100-lysed cells at the 36 hour time point to obtain normalised values plotted as “% Sytox green^+^ cells” on the Y-axis. Lines represent a sigmoidal, 4PL fit. **(C)** Parental and sg-*RIPK1* HT-29 cells expressing eGFP-V5-tagged human ZBP1 (wild type, isoform 1) were infected with HSV-1 ICP6^mutRHIM^ (MOI of 5) for the indicated time. ZBP1-eGFP-V5 was immunoprecipitated (IP) using GFP-Trap beads and input and IP samples were analysed by western blotting. **(D)** Parental (clone B9) and sg-*RIPK1* HT-29 cells expressing wild type (isoform 1) human ZBP1-eGFP-V5 were infected for 9 hours with HSV-1 ICP6^mutICP6^ (MOI of 5) and cell lysates were analysed by semi-denaturing detergent agarose gel electrophoresis (SDD-AGE). **(E)** Parental and sg-*RIPK1* HT-29 cells expressing human ZBP1-eGFP-V5 (wild type, isoform 1) were pre-treated with Nec-1s (5 µM) or GSK’840 (1 µM) for 30 min. and then infected with either HSV-1 ICP6^WT^ or HSV-1 ICP6^mutRHIM^ (MOI of 5) for 9 hours. ZBP1-eGFP-V5 was immunoprecipitated (IP) using GFP-Trap beads and input and IP samples were analysed by western blotting. **(F)** ZBP1-eGFP-V5 expressing HT-29 cells (clone B9) were treated with Nec-1s (5 µM) or GSK’840 (1 µM). Cells were then left uninfected (CTRL) or infected with HSV-1 ICP6^mutICP6^ (MOI of 5) for 9 hours. Cell lysates were analysed to by semi-denaturing detergent agarose gel electrophoresis (SDD-AGE).

Immunoprecipitation of ZBP1 after HSV-1 ICP6^mutRHIM^ infection showed that the presence of RIPK1 was required to activate RIPK3 in the ZBP1 signalling complex (Fig. 7C). SDD-AGE further showed that RIPK1 promoted the assembly of RIPK3 oligomers without affecting the oligomerisation of ZBP1 itself (Figs. 7D and S7F). RIPK1’s kinase activity was necessary to promote RIPK3 activation, but it was not required to recruit either RIPK1 or RIPK3 into the human ZBP1 signalling complex (Fig. 7E). Blocking RIPK3’s kinase activity with GSK’840 also did not prevent RIPK1 or RIPK3 recruitment to ZBP1 (Fig. 7D). While the recruitment of RIPK1 and RIPK3 into the ZBP1 signalling complex occurred independently of RIPK1’s enzymatic function, ZBP1-induced oligomerisation of both RIPK1 and RIPK3 required the kinase activity of RIPK1, but not that of RIPK3 (Fig. 7F).

Together, these data show that not only initial ZBP1 condensate formation (see Fig. S4C-E) but also the assembly of SDS-resistant ZBP1 oligomers occurred independently of RIPK1 and RIPK3 and their kinase activities, suggesting that ZBP1 oligomerisation represents an upstream event in necroptosis induction. The formation RIPK1/3 oligomers, however, fully depended on the enzymatic function of RIPK1, which is consistent with the observation that the kinase activity of RIPK1 controls the ordered assembly of a functional RIPK1/RIPK3 necrosome during TNFR1-induced necroptosis (Chen *et al*., 2022).

## Discussion

Together, our data support a two-step ZBP1 activation model: in a first step, the interaction of ZBP1’s Zα domains with Z-nucleic acids results in the local concentration of ZBP1 into condensates that are partially reversible in nature. This is followed by a second step involving the RHIM-mediated assembly of a solid state amyloidal signalling complex that activates RIPK1, RIPK3 and MLKL to induce necroptosis. In line with the first step of our model we find that a truncated ZBP1 protein containing only the Zα domains is efficient at forming condensates independently of the RHIMs, which is similar to a recent study (Xie *et al*, 2024). These assemblies remain partially fluid, suggesting that ZBP1 condensates are formed through liquid-liquid phase separation (LLPS) that evolve into gel-like state. Indeed, apart from their capacity to bind Z-nucleic acids, some -but not all-Zα domains induce LLPS, a process that is facilitated by binding to Z-RNA (Diallo *et al*, 2022). The Zα domain of ADAR1 and that present in the fish cypridinid herpesvirus protein ORF112 are particularly adept at undergoing LLPS. While ZBP1’s Zα1 or Zα2 domains in isolation were not able to phase separate or only weakly, the tandem Zα1-Zα2 configuration performed better, at least in the presence of a crowding agent polyethylene glycol (Diallo *et al*., 2022). This may explain our observation that the naturally occurring human ZBP1 isoform 2, which only contains Zα2, has reduced activity compared to isoform 1 containing both Zα domains. This is different from another study in which removal of the Zα1 domain had no measurable impact on the induction of HSV-1 ICP6^mutRHIM^-induced necroptosis, while the Zα2 domain was essential for human ZBP1 activation (Amusan *et al*., 2025). Variable contributions of the Zα1 domain and an essential function of the Zα2 domain were also reported for mouse ZBP1 in the context of viral infections (Guo *et al*., 2018; Koehler *et al*., 2021; Maelfait *et al*., 2017; Sridharan *et al*., 2017; Thapa *et al*., 2016; Yang *et al*., 2020) and in mouse models of autoinflammation (Jena *et al*, 2024; Jiao *et al*, 2020; Kesavardhana *et al*, 2020). The differences in the contributions of the Zα1 domain to ZBP1 activation may be attributed to the sensitivity of the experimental readout, the expression levels of ZBP1 and/or strength of the stimulus. Interestingly, Zα2 engages with Z-DNA in a different manner than the Zα1 domain and has low affinity for B-DNA (Ha *et al*., 2008; Kim *et al*, 2011a; Kim *et al*, 2011b; Schwartz *et al*., 2001). How the different properties of both Zα domains relate to the ability of ZBP1 to form condensates and to induce downstream signalling remains to be experimentally addressed.

Oxidative stress results in the accumulation of Z-RNA into stress granules (Yang *et al*., 2023), membranelles organelles containing stalled translation initiating complexes that form through LLPS of G3BP1 (Protter & Parker, 2016). Recruitment of ZBP1 into stress granules has been reported to induce necroptosis (Szczerba *et al*., 2023; Yang *et al*., 2023). As such, partitioning of ZBP1 into Z-RNA-containing stress granules may promote RHIM-mediated assembly into an amyloidal signalling complex. In the context of HSV-1 infection, however, we find that stress granule formation is not required to induce necroptosis, suggesting that concentration of ZBP1 in these organelles in not a universal activation mechanism. We report that both A- and Z-form dsRNA accumulate during HSV-1 infection. Interestingly, recognition of A-RNA by 2’-5’ oligoadenylate synthase-like (OASL) induces LLPS and recruits ZBP1 and RIPK3 into OASL condensates, a process that stimulates necroptosis of MCMV-infected mouse cells (Lee *et al*, 2023). It will be interesting to determine whether A-RNA binding to OASL synergises with Z-RNA recognition by ZBP1 to form condensates and to establish functional ZBP1 signalling complexes upon HSV-1 infection.

We propose that the second step in ZBP1’s activation process is the RHIM-mediated assembly of an amyloidal signalling complex. The amyloidal nature of ZBP1 oligomers is supported by their stability in the presence of denaturing agents and the solid state of optogenetically activated ‘RHIMs-only’ ZBP1 in our study, and by the formation of amyloidal fibrils by truncated recombinant ZBP1 proteins encompassing RHIM-A, -B and -C observed by others (Li *et al*., 2012; Steain *et al*, 2020; Xie *et al*., 2024). Similar to its mouse orthologue, our data and that of others show that the RHIM-A of human ZBP1 is essential for necroptosis induction by recruiting RIPK1 and RIPK3 (Amusan *et al*., 2025; Upton *et al*., 2012). Mutation of RHIM-A also renders ZBP1 condensates partially dynamic, suggesting that RHIM-A also contributes to ZBP1 amyloid formation. This likely results from a reduction in homotypic ZBP1 interactions rather than stabilising effects through heterotypic ZBP1-RIPK1 and/or ZBP1-RIPK3 interactions since both RIPK1 and RIPK3 are dispensable for stable ZBP1 oligomerisation.

Removal of RHIM-B, -C and the C-terminal tail renders mouse ZBP1 constitutively active (Koerner *et al*., 2024), suggesting that these domains exert inhibitory functions. Our data show, however, that RHIM-B and RHIM-C both positively contribute to RIPK1/3 recruitment and necroptosis induction indicating that constitutive activity of C-terminally truncated ZBP1 is due to loss of its disordered C-terminal tail or determined by species differences. Unlike RHIM-A, RHIM-B and -C do not seem to contribute to the formation of stable ZBP1 condensates. It is possible that mutation of RHIM-B or RHIM-C results in the formation of partially signalling incompetent, yet stable, ZBP1 oligomers. Similarly, the herpesviral RHIM-containing proteins ICP6 and M45 do not substantially interfere with ZBP1 oligomerisation, yet completely prevent RIPK1/3 recruitment to ZBP1 and necrosome formation. As such ICP6 and M45 may interfere with ZBP1 signalling by promoting the assembly of dysfunctional ZBP1-M45 or ZBP1-ICP6 hetero-amyloids (Pham *et al*, 2019). Future structural studies will determine how the three of RHIMs of ZBP1 contribute to the assembly of functional ZBP1 amyloids and downstream RIPK1/3 activation and how viral RHIM-containing proteins interfere with this process.

The relationship between RIPK1 and ZBP1 appears different in mouse and human systems. Firstly, mouse genetics show that RIPK1 prevents spontaneous induction of ZBP1-mediated necroptosis (Lin *et al*, 2016; Newton *et al*, 2016). Mechanistically, RIPK1 recruits CASP8 to the ZBP1 complex and cleaves RIPK1 to inactivate its kinase activity (Imai *et al*., 2024). In contrast, we find that in human cells loss of RIPK1 expression does not result in spontaneous necroptosis induction by ZBP1. This implicates that spontaneous ZBP1-mediated necroptosis may -unlike suggested by mouse studies-not underlie the development of human pathologies caused by *RIPK1* deficiencies (Cuchet-Lourenco *et al*, 2018; Li *et al*, 2019). Secondly, while necroptosis can proceed independently of RIPK1 in mouse cells (Upton *et al*., 2012), our data show that RIPK1 kinase activity is essential to induce necroptosis in human cells. This was also shown by a recent study using elegant ‘RHIM domain-swapping’ experiments demonstrating that the RHIM of mouse RIPK3 has a higher affinity for ZBP1 thereby bypassing the need for RIPK1 to induce necroptosis (Amusan *et al*., 2025). We now demonstrate that while the RIPK1 kinase activity is redundant for the recruitment of RIPK1 and RIPK3 to ZBP1, it is essential for the formation of stable RIPK1/3 oligomers downstream of ZBP1. This is in line with the essential role of RIPK1 autophosphorylation in the formation of a functional necrosome downstream of TNFR1 (Chen *et al*., 2022). Of note, these observations imply that RIPK1 kinase inhibitors may be effective against possible pathological contributions of ZBP1-induced necroptosis in human inflammatory pathologies such as cleavage-resistant RIPK1-induced autoinflammatory syndrome (Lalaoui *et al*, 2020; Tao *et al*, 2020) or inflammatory diseases caused by *CASP8* deficiencies (Chun *et al*, 2002; Lehle *et al*, 2019).

Finally, it will be interesting to test whether this two-step ZBP1 activation model holds true for ZBP1 signalling pathways other than necroptosis such as NF-κB or CASP8 activation in cell-intrinsic responses to viral infection, in setting of autoinflammation, or the response of tumour cells to cancer treatments.

**Correspondence and requests for materials** should be addressed to J.M.

## Acknowledgements

We are grateful to the VIB-UGent IRC Transgenic core facility and the VIB Flow and Bioimaging Cores for training, support and access to the instrument park. We are grateful to Sudan He for providing the ICP6 antibody, to Siddharth Balachandran for providing the IAV/PR8 strain, and Jiahuai Han for providing the HSV-1 ICP6^WT^ and HSV-1 ICP6^mutRHIM^ strains. J.N. and L.N. were supported by an FWO PhD fellowship (3S012819). Research in the J.M. lab was supported by an Odysseus II Grant (3G0H8618), EOS INFLADIS (3G0I5722), a junior research grant (G031022N) from the Research Foundation Flanders (FWO), a BOF project grant (BOF/24J/2023/159), a BOF starting grant (BOF/STA/202402/002) from Ghent University, and an ERC consolidator grant (ZIGNALLING, Proposal n° 101126114, 41U07824). Funded by the European Union. Views and opinions expressed are however those of the author(s) only and do not necessarily reflect those of the European Union or the European Research Council Executive Agency. Neither the European Union nor the granting authority can be held responsible for them. Research in the P.V lab was supported by FWO research projects (3G033120, 3G0A9322), EOS research consortium (EOS 30826052, 3G0I5722), UGent Special Research Fund Methusalem (01M00709), iBOF (01IB3920), grants from the Foundation against Cancer (365A08921), CRIG and GGIG consortia, and VIB. The D.L. lab was supported by an EOS INFLADIS (3G0I5722) grant from the Research Foundation Flanders (FWO).

## Author contributions

Conceptualisation: Josephine Nemegeer, Jonathan Maelfait. Investigation: Josephine Nemegeer, Aynur Sönmez, Evelien Dierick, Leslie Naesens, Denis L.J. Lafontaine, Jonathan Maelfait. Methodology: Josephine Nemegeer, Denis L.J. Lafontaine, Jonathan Maelfait. Formal analysis: Josephine Nemegeer, Denis L.J. Lafontaine, Jonathan Maelfait. Resources: Peter Vandenabeele, Denis L.J. Lafontaine, Jonathan Maelfait. Supervision: Jonathan Maelfait. Writing – original draft: Josephine Nemegeer, Jonathan Maelfait.

## Figure legends

**Supplementary Figure 1.**
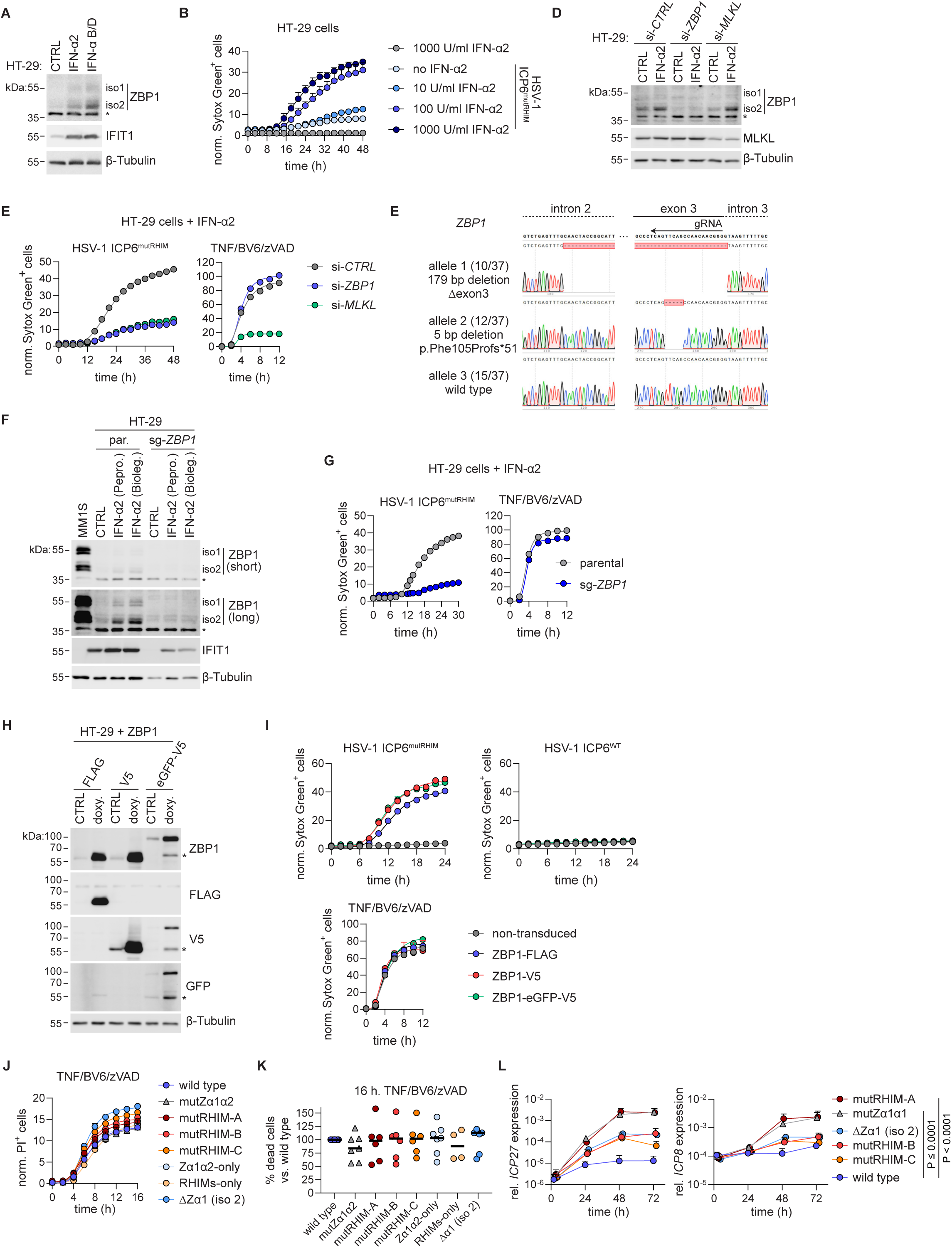
Endogenous human ZBP1 induces cell death and the Zα domains and RHIMs contribute to ZBP1-induced necroptosis. **(A)** HT-29 cells were treated with 1,000 U/ml of human IFN-α2 or recombinant IFN-α(B/D) hybrid for 18 hours and cell lysates were analysed by western blot. * represents a non-specific signal. **(B)** HT-29 cells were treated with indicated concentration of human IFN-α2 for 16 hours and subsequently infected with HSV-1 ICP6^mutRHIM^ at a multiplicity of infection (MOI) of 5. Cell death was quantified by measuring Sytox Green uptake every 2 hours using Incucyte cell imaging. The number of Sytox green^+^ cells per image at each time point was divided by the percentage of confluency to obtain normalised values plotted as “norm. Sytox green^+^ cells” on the Y-axis. Lines represent a sigmoidal, 4PL fit. **(C)** HT-29 cells were transfected with siRNAs targeting *ZBP1*, *MLKL* or a non-targeting control (si-*CTRL*) and 24 hours later cells were treated with 1,000 U/ml human. IFN-α2. 18 hours after IFN-α2 treatment, the cells were infected with HSV-1 ICP6^mutRHIM^ (MOI 5) or stimulated with 30 ng/ml TNF, 20 μM zVAD and 5 μM BV6. Cell death was measured by Sytox green uptake as in (B). **(D)** Western blot validation of siRNA-mediated knock-down for the experiment shown in (C). **(E)** *ZBP1* was targeted in HT-29 cells using CRISPR-Cas9-mediated gene editing. The genomic region targeted by the gRNA of the selected ZBP1-deficient HT-29 clone (sg-*ZBP1*) was PCR amplified, subcloned (n = 37), and analysed by Sanger sequencing. Two out of the three *ZBP1* alleles of HT-29 cells contained out of frame deletions resulting in deletion of exon 3 (Δexon3) or the introduction of a premature stop codon (p.Phe105Profs*51). One allele remained wild type. **(F)** Parental and sg-*ZBP1* HT-29 cells were incubated with 1,000 U/ml of two different commercial sources (Peprotech or Biolegend) of recombinant human IFN-α2 for 18 hours and cell lysates were analysed by western blotting. MM1S cells were used as a positive control for the ZBP1 antibody. **(G)** Parental and sg-*ZBP1* HT-29 cells were incubated with 1,000 U/ml human IFN-α2 for 18 hours and subsequently infected with HSV-1 ICP6^mutRHIM^ (MOI 5) or stimulated with 30 ng/ml TNF, 20 μM zVAD and 5 μM BV6. Cell death was measured by Sytox Green uptake as in (B). **(H)** HT-29 cells transduced with doxycycline-inducible lentivectors expressing C-terminally FLAG-, V5- or eGFP-V5-tagged wild type (isoform 1) human ZBP1 were treated or not with 1 µg/ml doxycycline for 24 hours and ZBP1 protein expression was analysed by western blotting. **(I)** HT-29 cells expressing FLAG-, V5- or eGFP-V5-tagged wild type (isoform 1) ZBP1 were infected with HSV-1 ICP6^mutRHIM^ (MOI of 5), HSV-1 ICP6^WT^ (MOI of 5) or stimulated with 30 ng/ml TNF, 20 μM zVAD and 5 μM BV6. Cell death was measured by Sytox green uptake as in (B). **(J)** HT-29 cells expressing the indicated eGFP-V5-tagged human ZBP1 variants were treated with 30 ng/ml TNF, 20 μM zVAD and 5 μM BV6. Cell death was measured by PI uptake as in Fig. 1B. **(K)** Percentage of norm. PI^+^ or Sytox green^+^ cells 16 hours after stimulation with necroptosis inducing cocktail as described in (L). Each data point represents an independent experiment. Values for wild type ZBP1 were set at 100 % within each experiment. **(L)** HT-29 cells expressing human ZBP1-eGFP-V5 variants were infected with HSV-1 ICP6^mutRHIM^ (MOI of 0.1). Viral replication was measured for indicated time points by determining relative mRNA expression of the HSV-1 *ICP27* and *ICP8* gene using RT-qPCR. P values by 2way ANOVA.

**Supplementary Figure 2.**
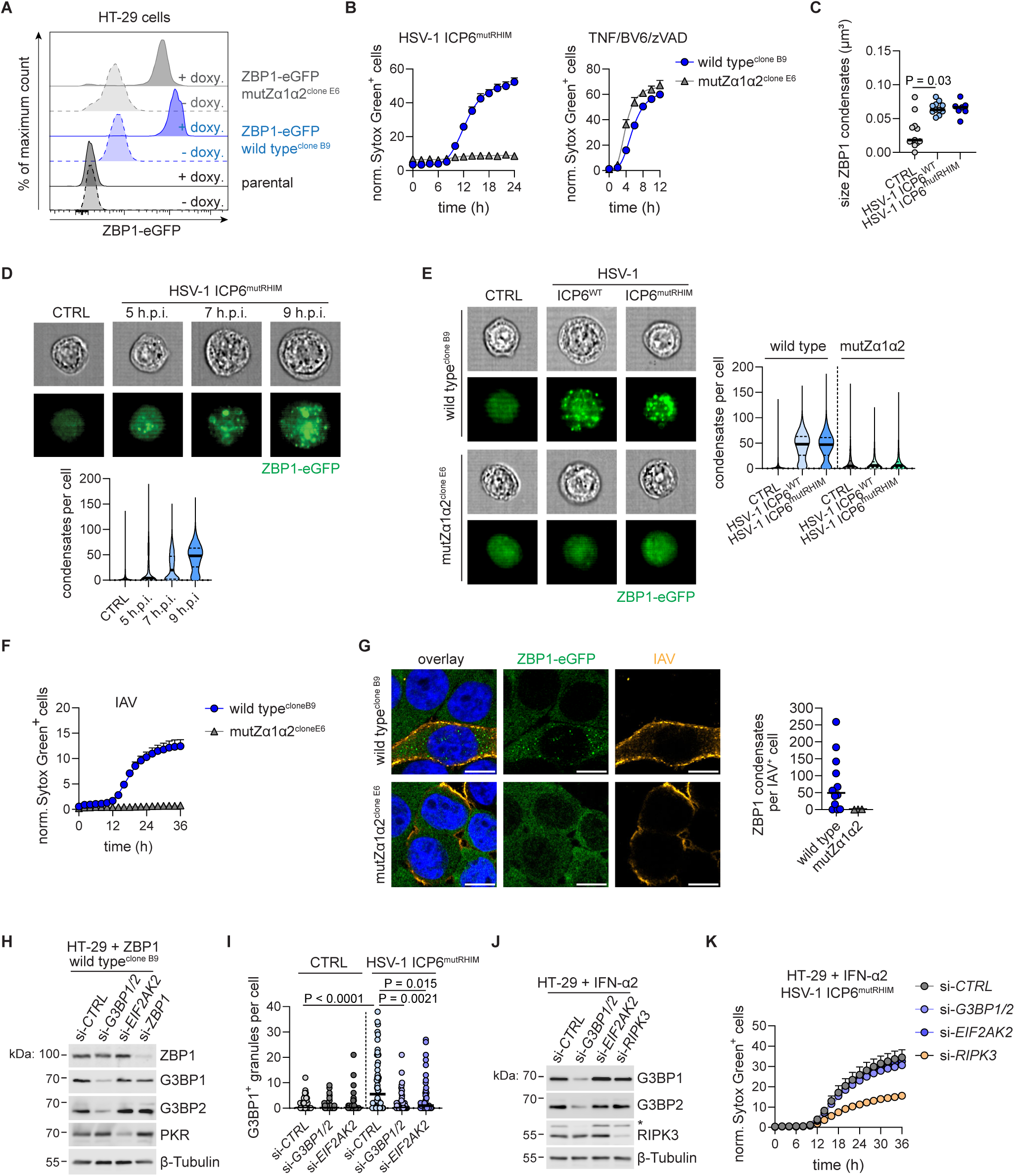
ZBP1 forms condensates after HSV-1 and IAV infection and ZBP1-dependent necroptosis occurs independently of stress granules. **(A)** Parental HT-29 cells and clones expressing doxycycline-inducible wild type (isoform 1, clone B9) or mutZα1α2 (clone E6) human ZBP1-eGFP-V5 were treated with 1 μg/ml doxycycline for 18 hours and ZBP1-eGFP expression was analysed by flow cytometry. **(B-E)** HT-29 cells expressing wild type (isoform 1) human ZBP1-eGFP-V5 (clone B9) or mutZα1Zα2 human ZBP1-eGFP-V5 (clone E6) were infected with HSV-1 ICP6^mutRHIM^ (MOI of 5) (B-E), HSV-1 ICP6^WT^ (C,E) or stimulated with 30 ng/ml TNF, 20 μM zVAD and 5 μM BV6 (B). **(B)** Cell death was quantified by measuring Sytox green uptake every 2 hours. The number of Sytox green^+^ cells per image at each time point was divided by the percentage of confluency to obtain normalised values plotted as “norm. Sytox green^+^ cells” on the Y-axis. Lines represent a sigmoidal, 4PL fit. **(C)** The size of wild type ZBP1-eGFP-V5 condensates was analysed using confocal microscopy. ZBP1 condensates were analysed in 3D-images and every dot represents a z-stack. P value by One-Way ANOVA. **(D,E)** Cells were analysed at the indicated time points after infection using imaging flow cytometry and ZBP1-eGFP condensates were quantified. Representative brightfield images and ZBP1-eGFP images are shown. **(F,G)** HT-29 cells expressing wild type (isoform 1) human ZBP1-eGFP-V5 (clone B9) or mutZα1Zα2 human ZBP1-eGFP-V5 (clone E6) were infected with Influenza A virus (IAV, MOI of 4). Scale bars, 5 µm. **(F)** Cell death was measured by Sytox Green uptake as in (B). **(G)** Representative images showing ZBP1-eGFP (green), DAPI (blue) and IAV proteins (yellow). Scale bar is 10 μm. ZBP1 condensates were analysed in 3D-images and every data point represents a z-stack of an infected cell. **(H,I)** HT-29 cells expressing wild type (isoform 1) human ZBP1-eGFP-V5 (clone B9) were transfected with siRNA targeting *G3BP1* and *G3BP2*, *EIF2AK2*, *ZBP1* or a non-targeting control (si-*CTRL*). 48 hours later, **(H)** Knock-down efficiency of was analysed by western blot or **(I)** cells were infected with HSV-1 ICP6^mutRHIM^ (MOI of 5) and the number of G3BP1^+^ granules were quantified per cell. Each dot represents a cell. P values by One-Way ANOVA. **(J,K)** HT-29 cells were treated with siRNA targeting *G3BP1* and *G3BP2*, *EIF2AK2*, *RIPK3* or a non-targeting control (si-*CTRL*). 24 hours later, cell were treated with 1,000 U/ml of human IFN-α2 and 18 hours later **(J)** Knock-down efficiency of was analysed by western blot (* represents the previously detected G3BP1 signal) or **(K)** cells were infected with HSV-1 ICP6^mutRHIM^ (MOI of 5) and cell death was measured by Sytox green uptake as in (B).

**Supplementary Figure 3.**
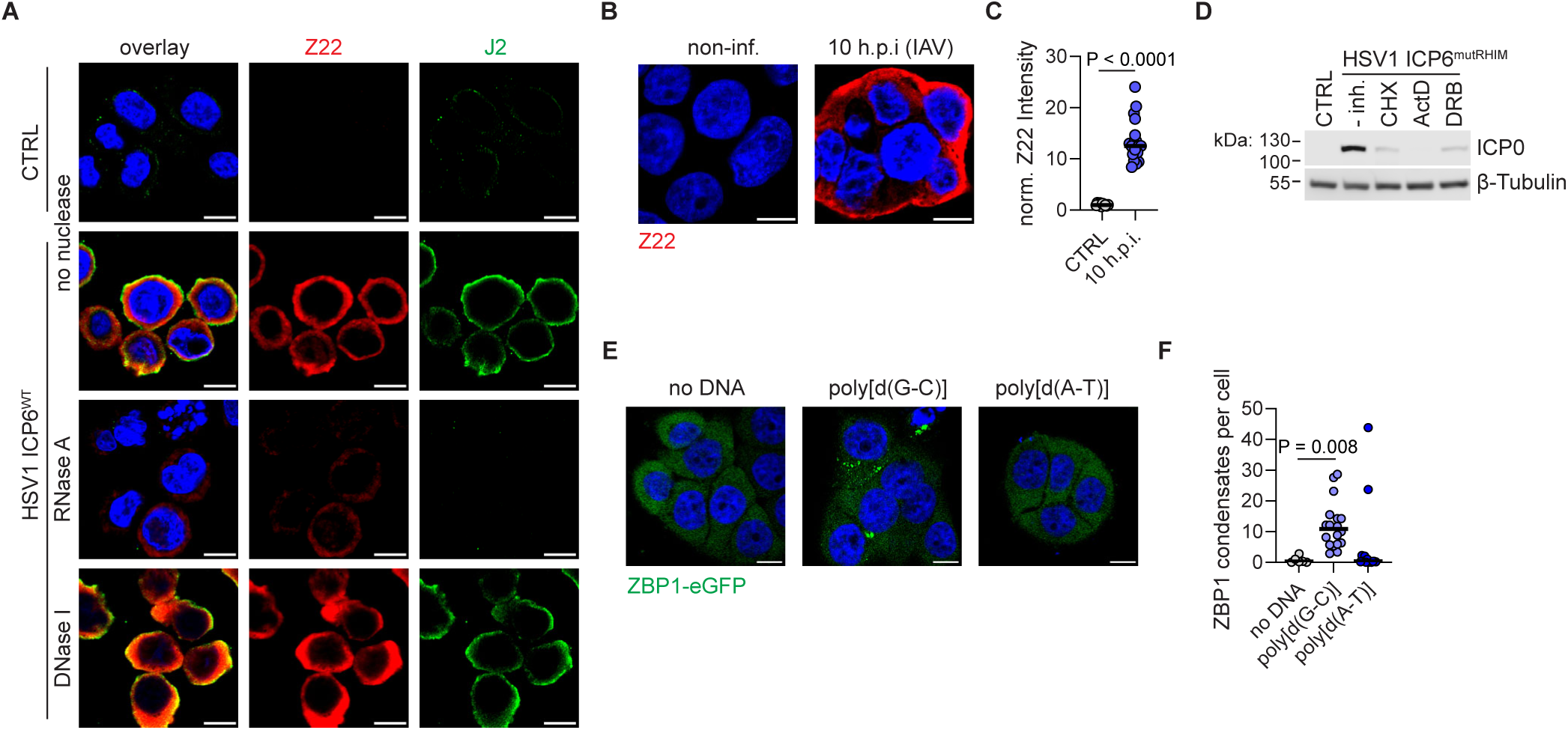
Z(-prone)-RNA accumulates in the cytosol after HSV-1 and IAV infection. **(A)** HT-29 cells were not infected (CTRL) or infected with HSV-1 ICP6^WT^ (MOI of 5) for 6 hours. Cells were treated with RNase A or DNase I before Z22 staining. Representative images show Z22 (red), an antibody recognising A-form dsRNA (J2, green) and DAPI (blue). Scale bars, 10 µm. **(B,C)** HT-29 cells were infected with IAV (MOI of 4) for 10 hours. **(B)** Representative images showing Z22 (red) and DAPI (blue). Scale bars, 10 µm. **(C)** Quantification of mean fluorescent intensity of Z22, normalised to the Z22 signal in non-infected (CTRL) cells, plotted as “norm. Z22 intensity” on the Y-axis. Each dot represents a single cell. P value by Mann-Whitney test. **(D)** HT-29 cells were left untreated (-inh.) or treated with 1 μg/ml cycloheximide (CHX), 5 μM Actinomycin D (ActD) or 50 μM RNA polymerase II inhibitor 5,6-dichlorobenzimidazole riboside (DRB) and then infected with HSV-1 ICP6^WT^ (MOI of 5) for 6 hours. The effects of these compounds on expression of the HSV-1 protein ICP0 was tested by western blotting. **(E,F)** HT-29 cells expressing wild type (isoform 1) human ZBP1-eGFP-V5 (clone B9) were transfected with poly[d(G-C)] or poly[d(A-T)] for 16 hours. **(E)** Representative images show ZBP1-eGFP (green) and DAPI (blue). Scale bars, 10 µm. **(F)** Quantification of the number of ZBP1-eGFP condensates per cell. ZBP1 condensates were analysed in 3D-images and every dot represents a z-stack. P value by One-Way ANOVA.

**Supplementary Figure 4.**
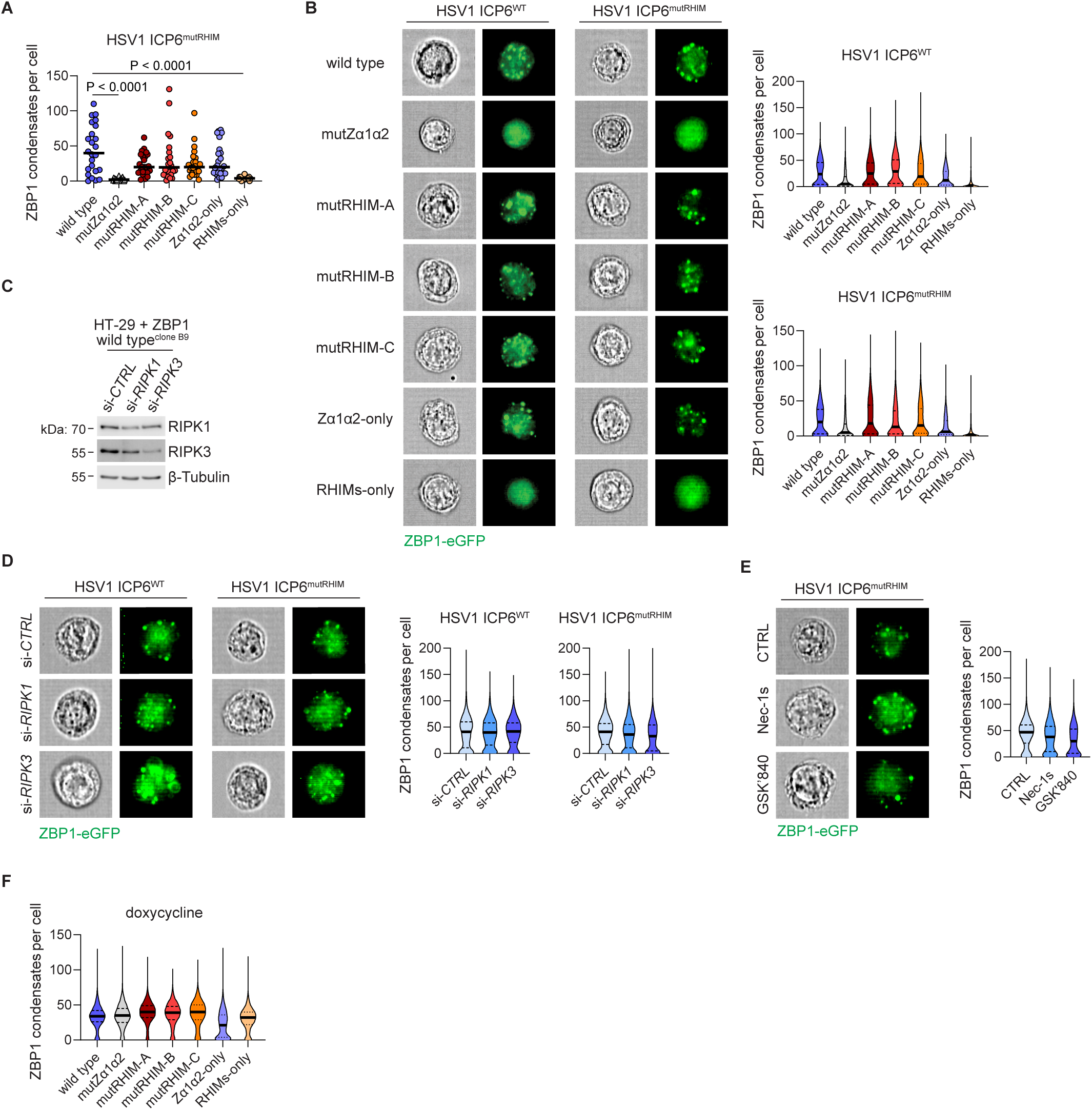
ZBP1 condensates form independently of the RHIMs and RIPK1/3. **(A)** HT-29 cells transduced with doxycycline-inducible lentivectors expressing the indicated human ZBP1-eGFP-V5 variants were infected with HSV-1 ICP6^mutRHIM^ at a multiplicity of infection (MOI) of 5. 9 hours post-infection cells were analysed by confocal microscopy. Quantification of the number of ZBP1 condensates per cell. ZBP1 condensates were analysed in 3D-images and every dot represents a single cell. P value by One-Way ANOVA. **(B)** HT-29 cells expressing the indicated human ZBP1-eGFP-V5 variants were infected with HSV-1 ICP6^WT^ or HSV-1 ICP6^mutRHIM^ (MOI of 5). 9 hours later, cells were analysed using imaging flow cytometry and ZBP1-eGFP condensates were quantified. Representative brightfield images and ZBP1-eGFP images are shown. **(C,D)** HT-29 cells expressing wild type (isoform 1) human ZBP1-eGFP-V5 (clone B9) were transfected with siRNAs targeting *RIPK1*, *RIPK3* or a non-targeting control (si-*CTRL*). 48 hour later, **(C)** Knockdown efficiency of was validated by western blotting or **(D)** cells were infected with HSV-1 ICP6^WT^ or HSV-1 ICP6^mutRHIM^ (MOI of 5). 9 hours later, cells were analysed as in (B). **(E)** HT-29 cells expressing wild type (isoform 1) human ZBP1-eGFP-V5 (clone B9) were pre-treated with Nec-1s (5 µM) or GSK’840 (1 µM) for 30 minutes and cells were then infected with HSV-1 ICP6^mutRHIM^ (MOI of 5). 9 hours later, cells were analysed as in (B). **(F)** HT-29 cells expressing the indicated doxycycline-inducible human ZBP1-eGFP-V5 variants were treated with 1 μg/ml doxycycline for 24 hours. Cells were analysed as in (B).

**Supplementary Figure 5.**
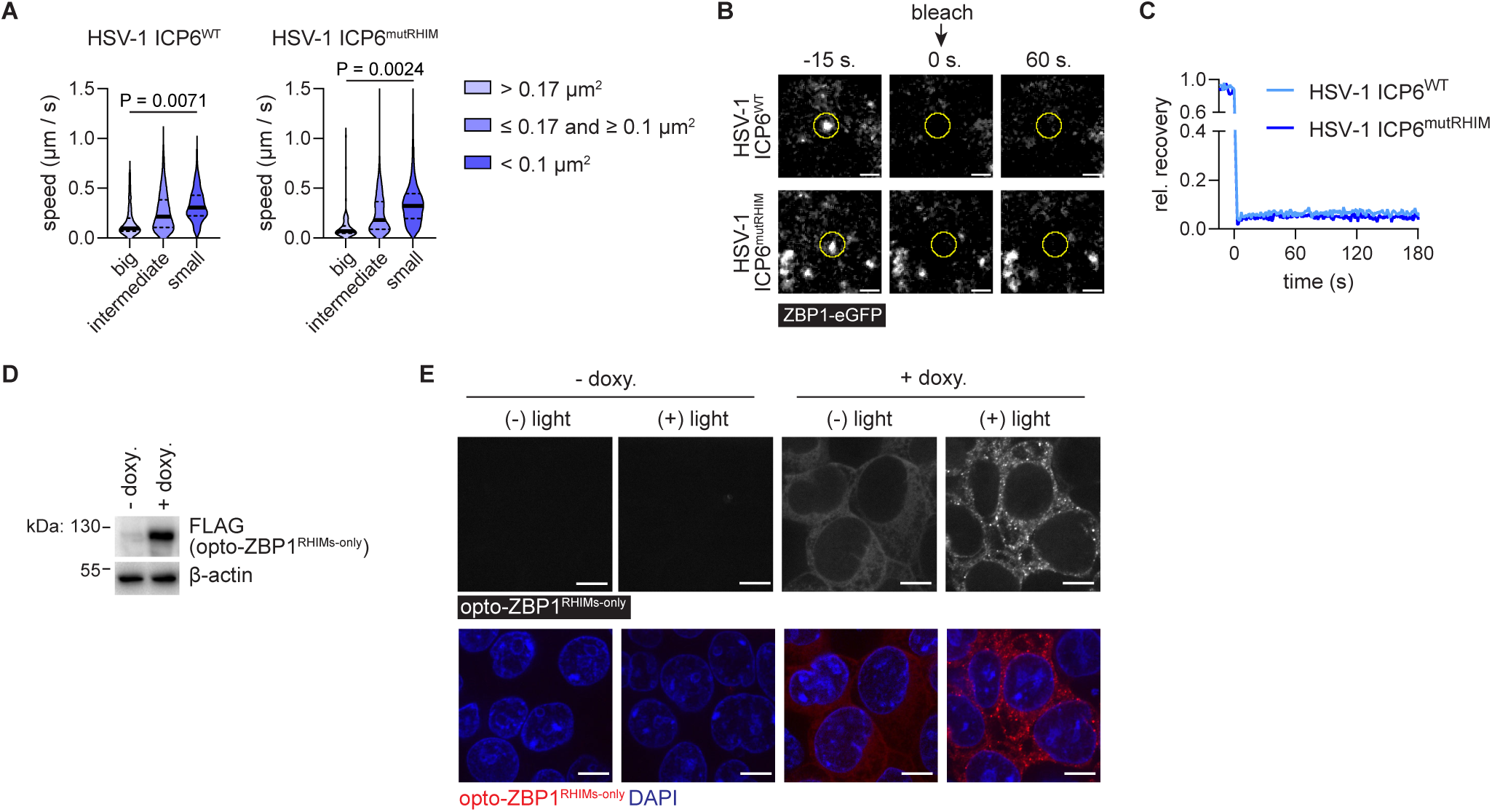
The RHIMs of ZBP1 support the formation of solid state condensates. (A-C) HT-29 cells expressing wild type (isoform 1) human ZBP1-eGFP-V5 (clone B9) were infected with HSV-1 ICP6^WT^ or HSV-1 ICP6^mutICP6^ (MOI of 5) for 8 hours. **(A)** The movement of condensates (n ≥ 92) of different sizes (big: > 0.17 um^2^; intermediate: ≤ 0.17 and ≥ 0.1 um^2^; small: < 0.1 um^2^;) were tracked over time and their respective speed (µm/s) was calculated. P values by One-Way ANOVA. **(B)** Representative images of ZBP1 condensates, before (-15 s.), immediately after (0 s.) or 60 seconds (60 s.) after photobleaching. The bleached areas are highlighted with a yellow circle. Scale bars, 2 µm. **(A)** Fluorescent recovery after photobleaching (FRAP) of ZBP1 condensates (n = 5) formed by the indicated human ZBP1 variants. The fluorescent intensity of the photobleached area at the indicated time point was normalised to the average fluorescent intensity at -15 s., which was set at 1, and plotted at as “rel. recovery” in the Y-axis. (**D**) Flp-In 293 T-REx cells expressing opto-ZBP1^RHIMs-only^ under a doxycycline-inducible promotor were left untreated (-doxy.) or treated with 1 µg/ml doxycycline (+ doxy.) for 24 hours and cell lysates were analysed by western blotting. **(E)** Opto-ZBP1^RHIMs-only^ expressing Flp-In 293 T-REx cells were left untreated (-doxy.) or treated with 1 µg/ml doxycycline (+ doxy.) for 24 hours. Cells were then kept in the dark [(-) light)] or exposed to 10 V blue light [(+) light)] for 2 min. Cells were fixed 10 min. after light exposure and analysed by confocal microscopy. Representative images showing opto-ZBP1^RHIMs-only^ (red) and DAPI (blue). Scale bars, 5 µm.

**Supplementary Figure 6.**
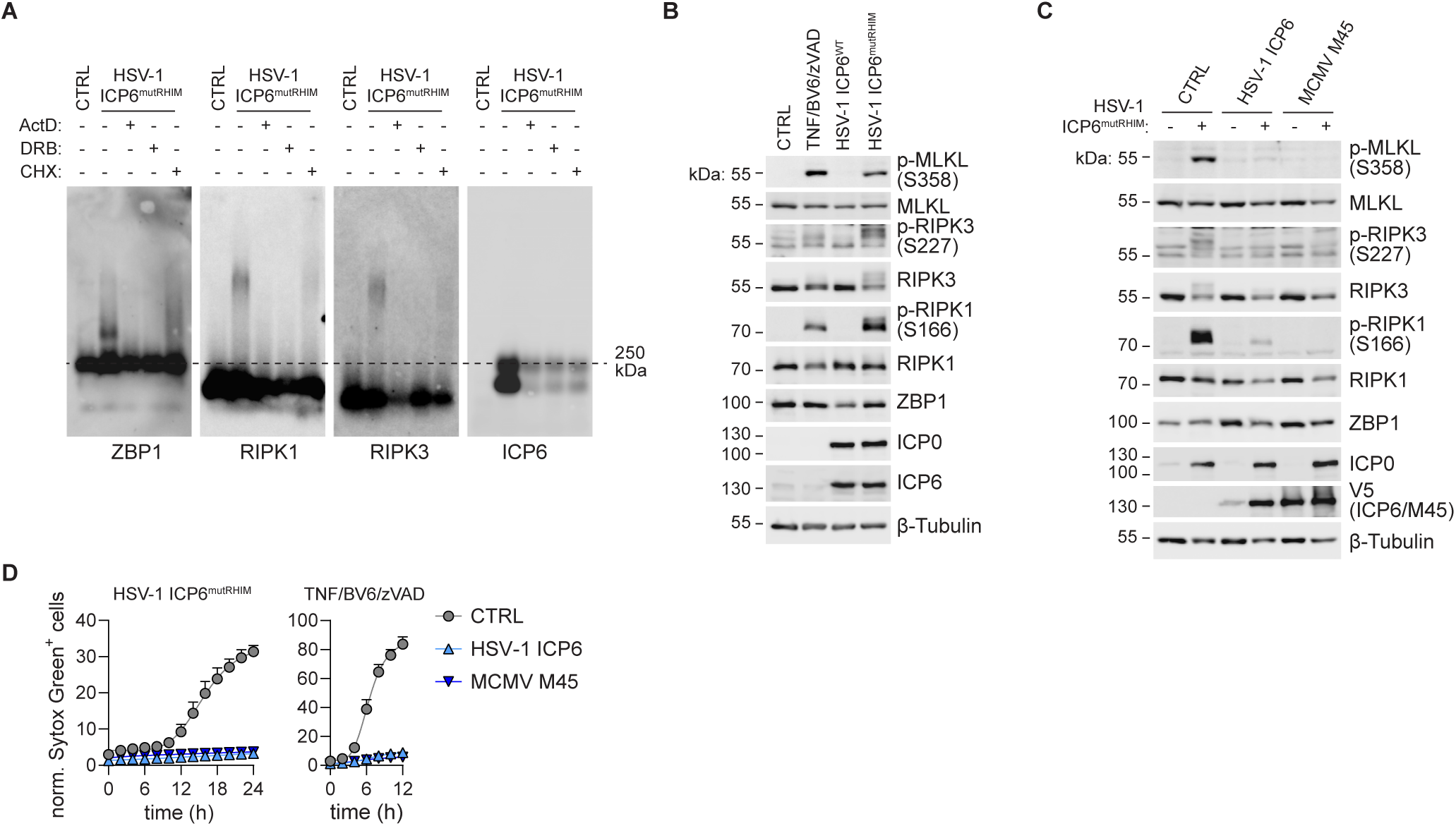
ZBP1 forms an amyloidal signalling complex independently of RIPK1/3 and ICP6 and M45 inhibit ZBP1-induced RIPK1/3 oligomerisation. **(A)** ZBP1-eGFP-V5 expressing HT-29 cells (clone B9) were treated with 5 μM actinomycin D (ActD), 50 μM RNA polymerase II inhibitor 5,6-dichlorobenzimidazole riboside (DRB) or 1 μg/ml cycloheximide (CHX). Cells were then left uninfected (CTRL) or infected with HSV-1 ICP6^mutICP6^ (MOI of 5) for 9 hours. Cell lysates were analysed by semi-denaturing detergent agarose gel electrophoresis (SDD-AGE). **(B)** HT-29 cells expressing wild type (isoform 1) human ZBP1-eGFP-V5 (clone B9) were infected with HSV-1 ICP6^WT^ or HSV-1 ICP6^mutICP6^ (MOI of 5) or stimulated with 30 ng/ml TNF, 20 μM zVAD and 5 μM BV6 and cell lysates were analysed by western blotting. **(C,D)** HT-29 cells expressing wild type (isoform 1) human ZBP1-eGFP-V5 (clone B9) were transduced with lentivectors expressing the viral RHIM-containing proteins HSV-1 ICP6 or MCMV M45 and the cells were infected with HSV-1 ICP6^mutRHIM^ (MOI of 5). **(C)** 9 hours later, cell lysates were analysed by western blotting. **(D)** Cell death was quantified by measuring Sytox green uptake every 2 hours using Incucyte cell imaging. The number of Sytox green^+^ cells per image at each time point was divided by the percentage of confluency to obtain normalised values plotted as “norm. PI^+^ cells” on the Y-axis. Lines represent a sigmoidal, 4PL fit.

**Supplementary Figure 7.**
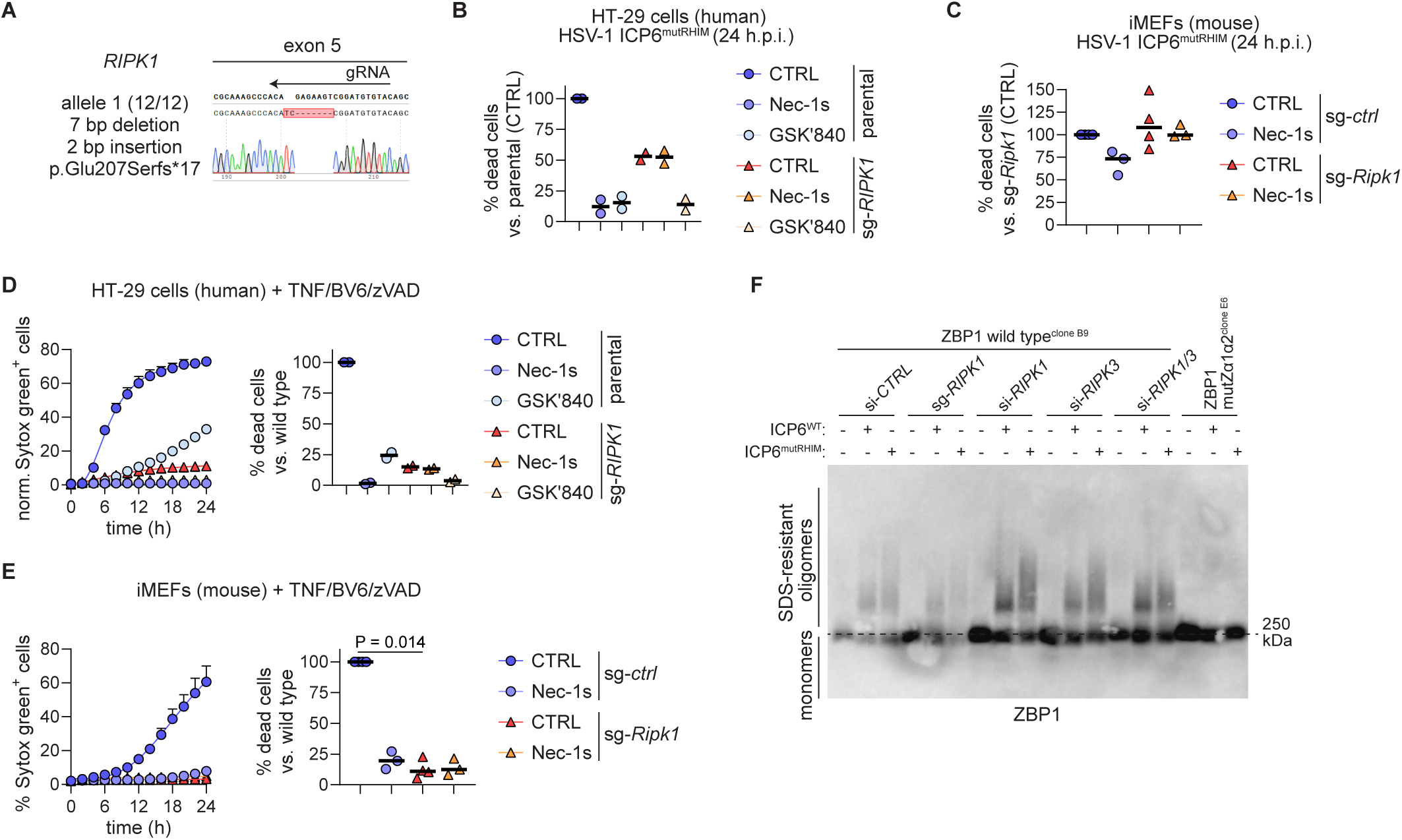
The kinase activity of RIPK1 is required for human ZBP1-induced necroptosis. **(A)** The genomic region targeted by the gRNA of the selected RIPK1-deficient HT-29 clone (sg-*RIPK1*) was PCR amplified, subcloned (n = 12), and analysed by Sanger sequencing. All *RIPK1* alleles of the clone contained out of frame mutations resulting in the introduction of a premature stop codon (p.Glu207Serfs*17). **(B)** Parental (clone B9) and sg-*RIPK1* HT-29 cells expressing human ZBP1-eGFP-V5 (wild type, isoform 1) were untreated (CTRL) or pre-treated with either Nec-1s (5 µM) or GSK’840 (1 µM) and then infected with HSV-1 ICP6^mutRHIM^ (MOI of 5). The graph shows the percentage of norm. Sytox green^+^ cells 24 hours after infection, calculated as described in Fig. 7A. Each data point represents an independent experiment. Values for parental cells that were not treated with RIPK1/3 inhibitors (CTRL) were set at 100 % within each experiment. **(C)** Sg-*ctrl* and sg-*Ripk1* immortalised mouse fibroblasts (iMEFs) were untreated (CTRL) or pre-treated with Nec-1s (5 µM) for 30 minutes and then infected with HSV-1 ICP6^mutRHIM^ (MOI of 5). The graph shows the percentage of norm. Sytox green^+^ cells 24 hours after infection as described in Fig. 7B. Each data point represents an independent experiment. Values for sg-*ctrl* cells that were not treated with the RIPK1 inhibitor (CTRL) were set at 100 % within each experiment. **(D)** Parental (clone B9) and sg-*RIPK1* HT-29 cells expressing human ZBP1-eGFP-V5 (wild type, isoform 1) were pre-treated for 30 minutes with either Nec-1s (5 µM) or GSK’840 (1 µM) and then stimulated with 30 ng/ml TNF, 20 μM zVAD and 5 μM BV6. Cell death was measured by Sytox green uptake. Left graph: the number of Sytox green^+^ cells was analysed as in Fig. 7A. Right graph: percentage of norm. Sytox green^+^ cells 16 hours after stimulation as in (B). **(E)** Sg-*ctrl* and sg-*Ripk1* immortalised mouse fibroblasts (iMEFs) were pre-treated with Nec-1s (5 µM) for 30 minutes and then and then stimulated with 30 ng/ml TNF, 20 μM zVAD and 5 μM BV6. Cell death was measured by Sytox green uptake. Left graph: the number of Sytox green^+^ cells was analysed as in Fig. 7B. Right graph: percentage of norm. Sytox green^+^ cells 16 hours after stimulation as in (C). **(F)** HT-29 clones expressing wild type (isoform 1, clone B9) were transfected with siRNA targeting *RIPK1*, *RIPK3*, *RIPK1* and *RIPK3*, or a non-targeting control (si-*CTRL*) for 48 hours. Sg-*RIPK1* wild type (isoform 1) human ZBP1-eGFP-V5 and mutZα1Zα2 human ZBP1-eGFP-V5 expressing HT-29 cells were included as controls. Cells were infected with either HSV-1 ICP6^WT^ or HSV-1 ICP6^mutICP6^ (MOI of 5) for 9 hours and cell lysates were analysed by SDD-AGE.

## Movie legends

### Movie 1. ZBP1 forms condensates and induces necroptosis after ICP6 RHIM-mutant HSV-1 infection

Live cell imaging of HT-29 cells expressing wild type (isoform 1) human ZBP1-eGFP-V5 (clone B9) infected with HSV-1 ICP6^mutICP6^ (MOI of 5). Propidium iodide (PI) was added to the medium to visualise cells that have lost plasma membrane integrity. Scale bar, 20 µm. Timestamp scale, hours (H).

### Movie 2. ZBP1 condensate formation and necroptosis induction requires intact Zα domains

Live cell imaging of HT-29 cells expressing Zα domains mutant (mutZα1α2) human ZBP1-eGFP-V5 (clone E6) infected with HSV-1 ICP6^mutICP6^ (MOI of 5). Propidium iodide (PI) was added to the medium to visualise cells that have lost plasma membrane integrity. Scale bar, 20 µm. Timestamp scale, hours (H).

### Movie 3. ICP6 does not prevent ZBP1 condensate formation, but inhibits necroptosis induction

Live cell imaging of HT-29 cells expressing wild type (isoform 1) human ZBP1-eGFP-V5 (clone B9) infected with HSV-1 ICP6^WT^ (MOI of 5). Propidium iodide (PI) was added to the medium to visualise cells that have lost plasma membrane integrity. Scale bar, 20 µm. Timestamp scale, hours (H).

### Movies 4 and 5. ZBP1 assembles into solid state foci and ICP6 does not change this process

Live cell imaging of ZBP1-eGFP fluorescence recovery after photobleaching. ZBP1 condensate formation was induced by infecting wild type (isoform 1) human ZBP1-eGFP-V5 expressing HT-29 cells (clone B9) with HSV-1 ICP6^WT^ (movie 4) or HSV-1 ICP6^mutICP6^ (movie 5) (MOI of 5). Cells that contained large ZBP1-eGFP condensates were selected for FRAP analysis. Images were acquired over a 3 min. period at 1 second (s) intervals. The bleached area is indicated by a yellow circle. Scale bar, 5 µm. Timestamp scale, seconds (s). No fluorescence recovery was observed, indicating a solid material state.

### Movies 6 until 11. The RHIMs of ZBP1 are required to form solid state condensates

Live cell imaging of ZBP1-eGFP fluorescence recovery after photobleaching. ZBP1 condensate formation was induced by infecting HT-29 cells expressing the indicated GFP-V5-tagged ZBP1 variants with HSV-1 ICP6^WT^ (MOI of 5). Cells that contained large ZBP1-eGFP condensates were selected for FRAP analysis. Images were acquired over a 3 min. period at 1 second (s) intervals. The bleached area is indicated by a coloured circle. Scale bar, 5 µm. Timestamp scale, seconds (s). No fluorescence recovery was observed except in ZBP1 variants in which RHIM-A was mutated or when the RHIMs were removed, indicating that the RHIMs contribute to the formation of solid state ZBP1 condensates.

### Movie 12. Kinetics of opto-ZBP1^RHIMs-only^ foci formation after a blue light pulse

Live cell imaging of opto-ZBP1^RHIMs-only^ foci formation. Expression of the construct was induced by treating cells with 1 µg/ml doxycycline for 24 hours. Cells were exposed to a 2 min., 10V blue light pulse and imaged every 30 s. for 1 hour. Scale bar, 5 µm. Timestamp scale, min. The movie shows the progressive clustering of opto-ZBP1^RHIMs-only^ into cytoplasmic foci. Initially, the foci are small and highly mobile, rapidly roaming through the cytoplasmic space. The first sizeable structures become visible approximately 10 min. after the blue light pulse. At later time points (∼45 min.), the foci coalesce into larger, less mobile aggregates.

### Movie 13. FRAP analysis of a spontaneously formed opto-ZBP1^RHIMs-only^ focus

Live cell imaging of opto-ZBP1^RHIMs-only^ fluorescence recovery after photobleaching. Opto-ZBP1^RHIMs-only^ expression was induced by treating cells with 1 µg/ml doxycycline for 27 hours. Cells that contained spontaneously formed foci were selected for FRAP analysis. Images were acquired over a 5 min. period at 5 s. intervals. The bleached area is indicated by a dotted circle. Scale bar, 2 µm. Timestamp scale, seconds (s.). No fluorescence recovery was observed, indicating a solid material state.

### Movie 14. FRAP analysis of a blue light-induced opto-ZBP1^RHIMs-only^ focus

Live cell imaging of opto-ZBP1^RHIMs-only^ fluorescence recovery after photobleaching. Opto-ZBP1^RHIMs-only^ expression was induced by treating cells with 1 µg/ml doxycycline for 24 hours. Cells were then exposed to a 2 min., 10 V blue light pulse, and FRAP analysis was performed 3 hours later. Images were acquired over a 5 min period at 5 s intervals. The bleached area is indicated by a dotted circle. Scale bar, 2 µm. Timestamp scale, seconds (s.). No fluorescence recovery was observed, indicating a solid material state.

## Methods

### Cell culture

HEK293T cells, immortalised mouse embryonic fibroblasts (iMEF), Flp-In 293 T-REx and Vero cells (ATCC) were maintained in Dulbecco’s modified Eagle’s medium (DMEM), high glucose (Gibco, 11965092). HT-29 cells were kept in McCoy’s 5A medium (Gibco, 16600082). Media contained high glucose were supplemented with 10 % foetal bovine serum (Gibco or Tico), 1 mM sodium pyruvate (Sigma-Aldrich, S8636) and 2 mM L-glutamine (Sigma-Aldrich, G7513). All cells were maintained at 37°C with 5% carbon dioxide.

### Production of transduced cell lines

Cell lines, stably expressing a protein of interest, were made using lentiviral vectors. Lentiviral vectors were made in HEK293T cells, transfected with C-terminal eGFP/V5-, V5- or FLAG-tagged human ZBP1 variants transducing vectors in the pDG2i backbone together with the pCMV delta R8.91 gag-pol-expressing packaging plasmids and pMD2.G VSV-G-expressing envelope plasmid. Included ZBP1 variants were wild type ZBP1 (iso1), the natural splice variant lacking the first Zα domain [ΔZα1 (iso 2)], a Zα1α2 mutant (N46A/Y50A and N141A/Y145A), separate RHIM mutants [^205^IQIG>AAAA (mutRHIM-A), ^264^VQLG>AAAA (mutRHIM-B) or ^332^ATIG>AAAA (mutRHIM-C)], Zα1α2 only (amino acids 1-169) and the RHIM only (amino acids 193-429). In brief, HEK293T cells were reverse transfected with the mix of plasmids in a 6 well plate, using approximately 750,000 cells per well. All transfections were done using Lipofectamine 2000 (Invitrogen, 11668-027) in a 2:1 ratio (2 µl Lipofectamine per 1 µg of DNA). 24 hours after transfection medium was refreshed. The lentiviral vector containing supernatants was collected 48 hours later, passed through a 0.45 µm filter (Thermofisher, Merck Millex, SLHVR33RB) and frozen at -80°C until transduction. Lentiviral transduction of HT-29 cells was done in a 6-well by spin-fection (1 hour, 800g, 32°C) in the presence of polybrene (8 µg/ml, Sigma-Aldrich, TR-1003-G). Transduced cells were selected using either blasticidin S (10 µg/ml, Invitrogen, R210-01) or puromycin (1 µg/ml, Sigma-Aldrich, P-7255) for 2 weeks. Polyclonal cell lines expressing eGFP/V5-tagged ZBP1 variants expressing equivalent protein levels were made by cell sorting with BD FACS Melody cell sorter (Biosciences) using a narrow margin on the eGFP-signal. For microscopy and image stream purposes clonal cell lines expressing either wild type or mutZα1Zα2 ZBP1 were produced by single cell sorting. Clonal cell lines were validated afterwards to ensure an equal expression level via flow cytometry and western blotting.

### Production of knock-out cell lines

HT-29 knock-out cell lines were produced via electroporation of Cas9-RNPs. In brief, 0.2 nmol crRNA (IDT) and 0.2 nmol tracrRNA (IDT) were mixed, denatured for 5 minutes at 95°C and annealed for 20 minutes at room temperature. Next, the RNA duplex was combined with 20 µg GFP-tagged Cas9 (VIB Protein Service Facility) and incubated for 10 minutes at room temperature. Finally, Cas9-RNPs were combined with 1.25 x 10^6^ cells and 0.2 nmol Electroporation enhancer (1075915, IDT) in 100 µl. Electroporation was done using the NEPA21 electroporator (NepaGene). In non-transduced HT-29 cells, electroporation was done with unlabelled tracrRNA (#1072533, IDT) and a GFP-tagged Cas9 (VIB Protein Service Facility), targeting ZBP1 (5’-CCCGTTGTTGGCTGAACTGA-3’, IDT). Electroporation of HT-29 cells expressing eGFP/V5-tagged ZBP1 was done with ATTO^TM^ 647-labelled tracrRNA (#10007853, IDT) and unlabelled Cas9 (VIB Protein Service Facility) targeting RIPK1 (5’-TACACATCCGACTTCTCTGT-3’, IDT). Sixteen hours after electroporation, single GFP or ATTO^TM^ 647 positive cells were sorted (Melody, Biosciences) and plated in 96 well plate. Cell lines were screened via PCR and hits were validated with western blotting and subcloning. Subcloning was done using Zero Blunt TOPO PCR Cloning Kit (Invitrogen; 450245). At least 12 subclones were individually sequenced.

To produce RIPK1 knock-out iMEF cells LentiCRISPRv2-generated lentiviral vectors were used. Summarised, lentiviral vectors were produced in HEK293T cells by co-transfection of psPAX2, p-CMV-VSV-G and lentiCRISPRv2 plasmids targeting RIPK1 (5’-CCTGAATTTGACCTGCTCGG-3’, IDT) in HEK293T cells via Calcium Phosphate transfection. Lentiviral vector supernatant was harvested 48h following transfection and subsequently used to transduce iMEFs in the presence of 8 µg/mL Polybrene (H9268, Sigma-Aldrich). The next day, transduced cells were selected with 2 µg/mL Puromycin (P-7255, Sigma-Aldrich) for the duration of one week. Efficiency of RIPK1 knock-out in polyclonal MEF cells was validated using western blotting and cell death assays.

### Generation of Cry2olig-mCherry-ZBP1 cell lines

The opto-ZBP1 plasmid (pDL1143) was generated as follows: the pcDNA5-FRT/TO plasmid was linearized with HindIII and XhoI, and a custom gBlock (IDT) containing the truncated ZBP1 sequence was inserted using InFusion cloning. The resulting plasmid was then linearized using BamHI and SbfI, and an insert amplified from a Cry2olig-mCherry-containing plasmid (Addgene) was inserted using InFusion cloning. The opto-ZBP1 cell line (clone #1) was generated as follow: HEK293 Flp-In™ T Rex™ cells were co-transfected with pDL1143 and pOG44 (encoding the recombinase) according to the manufacturer’s instructions. Integration of opto-ZBP1 at the FRT locus was selected using hygromycin.

### siRNA-mediated knockdown

Transient knockdown of *RIPK1*, *RIPK3*, *MLKL*, *G3BP1*, *G3BP2*, *PKR* or *ZBP1* was achieved via reverse transfection of siRNA targeting *RIPK1* (ON-TARGETplus, SMARTpool L-004445-00-0005, Dharmacon), *RIPK3* (Accell, SMARTpool E-003534-00-0005, Dharmacon), *MLKL* (ON-TARGETplus, SMARTpool L-005326-00-0005, Dharmacon), *G3BP1* (ON-TARGETplus, SMARTpool L-012099-00-0005, Dharmacon), *G3BP2* (ON-TARGETplus, SMARTpool L-015329-01-0005, Dharmacon), *PKR* (ON-TARGETplus, SMARTpool L-003527-00, Dharmacon) or *ZBP1* (ON-TARGETplus, SMARTpool L-014650-00-0005, Dharmacon). As a control for baseline cellular responses to siRNA, a non-targeting pool was transfected (ON-TARGETplus, Non-targeting Control Pool D-001810-10-20, Dharmacon). Transfections were done with DharmaFECT-1 (Dharmacon, T-2001-03) following manufacturer’s instructions. Assays were performed 48 hours post transfection with siRNA. Knockdown efficiency was validated using qPCR targeting downregulated gene and/or by following protein abundance via western blotting.

### Viruses and infection protocol

HSV-1 viruses encoding either a wild-type (WT) ICP6 (HSV-1 ICP6^WT^) or an ICP6 RHIM mutant (HSV-1 ICP6^mutRHIM^) were made by dr. Jiahuai Han (Xia Men University, Xiamen, China) (Huang *et al*., 2015), and kindly provided by prof. William J. Kaiser (Emory Vaccine center, Emory University, Atlanta, USA). HSV-1 viruses were propagated in Vero cells. The cells were inoculated with a multiplicity of infection (MOI) of 0.01 for 2 hours in serum-free DMEM, supplemented with sodium pyruvate (Sigma-Aldrich, S8636) and L-glutamine (Sigma-Aldrich, G7513). Virus was harvested after 48 hours, when 100% cytopathic effect (CPE) was reached. Next, the Vero cells were released using cell scrapers (Cole-Parmer, # WZ-04396-54) and the medium containing both cells and virus was spun down at 1,200 g for 5 minutes at 4 degrees Celsius. The supernatant was collected and the remaining cells were disrupted via repeated freeze-thaw cycles. After a second spin of the cells (1,700 g, 5 minutes, 4°C), all supernatant was collected with careful consideration not the disrupt the cell pellet. Supernatants containing HSV-1 particles was then concentrated by ultracentrifugation (20,000 RPM, 1 h, 4°C, SS34 rotor) and stored in serum-free DMEM, supplemented with 10% glycerol at -80°C. Viral titres were quantified using a standard plaque assay on Vero cells. In brief, Vero cells were infected with a dilution series of the virus stock for 2 hours in serum-free medium. Afterwards, virus-containing medium was washed away and replaced by a semisolid matrix (full strength DMEM + 1.5% carboxymethyl cellulose (CMC)). After 2 days, cells were washed and fixed with 4% PFA (SANBIO, AR1068) for 30 minutes and subsequently stained with Crystal violet (Sigma-Aldrich, V5265) at room temperature for 3 minutes. After thoroughly washing with distilled water, the plates were airdried and quantified. The dilution series and quantification was always done in duplicate.

IAV PR/8 virus was kindly provided by prof. Siddharth Balachandran (Blood Cell Development and Function Program, Fox Chase Cancer Center, Philadelphia, PA). Cells were washed cells with serum-free medium and subsequently infected with IAV in serum-fee medium for 1 hour at 37°C. Next, the virus-containing medium was removed and interchanged for serum-containing medium.

### DNA transfection

HT-29 cells were seeded in an 8-well microscopy chamber (iBidi, 80806), using 90 000 cells per well. After 24 hours cells were transfected with 500 ng of poly(dC:dG):poly(dG:dC) (Invivogen, tlrl-pgcn) or poly(dA:dT):poly(dT:dA) (Invivogen, tlrl-patn) using Lipofectamine 2000 (Invitrogen, 11668-027) in a 1:3 ratio, following manufacturer’s instructions. Next, cells were left for 8 hours and fixed, as described in ‘confocal microscopy’. For plasmid transfections HEK cells were reverse transfected with tagged human or mouse RHIM proteins using lipofectamine in a 1:2 ratio, following manufacturer’s instructions. Cells were left for 24 hours and processed, as described in ‘co-immunoprecipitation’.

### Confocal microscopy

For all confocal microscopy experiments, cells were seeded into an 8-well coverslip (iBidi, 80826). Cells were fixed with 4% PFA (SANBIO, AR1068) for 30 minutes at room temperature. Next, cells were washed thoroughly with PBS and permeabilized with 0.5% Triton X-100 (Sigma Aldrich, 9036-19-5) in PBS for 30 minutes. The coverslip was then blocked for 2 hours at room temperature with Maxblock (Active Motif, 15252). Subsequently, primary antibodies, including mouse anti-ICP0 (Santa Cruz, Sc-53070, 1/50), rabbit anti-G3BP1 (Cell Signaling, #61559, 1/200), mouse polyclonale anti-IAV (Produced in-house, kindly provided by the lab of Prof. X. Saelens, 1/100), rabbit anti-Z-DNA clone Z22 (Absolute antibodies, Ab00783-23-0, 1/200), mouse anti-dsRNA clone J2 (SCICONS, 10010200, 1/200) were incubated overnight at 4°C in 0.1% Triton X-100 in PBS. After three 5-minute wash steps with 0.1% Triton X-100, sample was incubated with secondary antibodies, including Goat anti-mouse DyLight 633 (Thermofisher, 35513, 1/1000), Goat anti-mouse DyLight 488 (Thermofisher, 35503, 1/1000), and DAPI (Thermofisher, D21490) in 0.1% Triton X-100 in PBS, shielded from light. Lastly, cell were washed repeatedly with PBS and stored in an excess PBS until imaging on the LSM880 confocal microscope (Zeiss). For the visualisation of Z-nucleic acids, the protocol was adapted to include a tyramide amplification step, as described in Nemegeer et. al JoVE (2022, DOI: 10.3791/64332-v). In short, the coverslip was treated with HRP-labelled anti-Rabbit antibody (ECL Anti-Rabbit IgG HRP, VWR, K4002) for 30 minutes after overnight incubation with primary antibodies. Then, sample was treated with biotinylated-tyramide (R&D systems, 6241) for 10 minutes after which the amplification was visualised using fluorophore-labelled streptavidin (Thermofisher, S11226, 1/500) for 2 hours before imaging together with secondary antibodies mix in normal staining protocol.

For live cell imaging, media of cells was supplemented with Hoechst 33342 (1/5000, Thermofisher, H3570) and Propidium Iodide (1/1000, Sigma-Aldrich, P-4170), 30 minutes prior to imaging. Imaging was done using the Spinning disk confocal microscope (Zeiss). Z-stacks were taken every 15 minutes. Data was processed using Image J (FIJI). Movies are represented as an extended depth of focus. Live aggregate tracking was done using the LSM880 confocal microscope (Zeiss).

### Confocal image processing and image analysis

Images made using the Fast Airyscan LSM880 confocal microscope (Zeiss), were processed using Airyscan processing (Zen black software, Zeiss). All represented images represent 1 z-dimension of the 3D image and were exported using Zen blue (Zeiss) or Image J (FIJI). **Quantification of ZBP1-GFP aggregates** after viral infection was done with Volocity 6.3 (Volocity). 3D images were loaded in a velocity library and represented in extended dept of focus. For aggregate quantification, aggregates were defined as > 0.01 µm³, < 10 µm³. To quantify the relative amount of aggregates per cell, the nuclei were counted using a threshold in size of > 150 µm³. In IAV infection assays, a marker for IAV infection (mouse polyclonal IAV antibody) was used to identity infected cells. Exported data was further processed with Excel. **Analysis of RNA/DNA accumulation** with Z22/J2 staining was done using the Arivis software (Zeiss). Cells were identified using a deep learning-based tool imbedded in the software. No threshold was used, median fluorescence intensity (MFI) was identified on a per cell basis. For **aggregate tracking** consecutive confocal images were made of infected cells and analysed with Arivis software (Zeiss). Aggregates were defined using the ‘Blob Finder’ feature, with a guideline diameter of 0.4 µm, a probability threshold of 8% and a split sensitivity of 90%. Next, aggregates were subdivided based on size, using ‘Object feature filter’ feature, to distinguish between small (>0.02 µm³, <0.1µm³) intermediate (>0.1µm³, <0.17 µm³) and big aggregates (>0.17µm³). These objects were tracked using ‘Brownian motion’ settings, with a maximum distance of 900 nm. Tracks were included if the aggregate could be followed for at least 3 consecutive images. Data, considering track speed (µm/s) and track length (µm), were further processed using Excel.

### Fluorescent Recovery After Photobleaching (FRAP)

HT-29 cells, expressing a eGFP-tagged ZBP1 variant, were infected with HSV-1 and visualized with LSM880 confocal microscope (Zeiss). The eGFP-positive aggregate was measured for 15s prior to bleaching, with images every second. Next, the region of interest was bleached using the 488 laser, with a laser power of 70 %, for 10 iterations and a scan-speed of 3. Recovery of bleached aggregate was followed over 3 minutes post bleaching, with images every second. To visualise aggregates, the pinhole of the microscope was set to 106 µm. Data analysis was done with Image J (Fiji), using Stowers ImageJ plugins. An individual spectrum was created (default, Avg) for each bleached aggregate, these were then combined and normalized (MIN/MAX settings). X/Y values were then exported and further processed in excel. Representative images were exported using Zen blue (Zeiss).

### Optogenetics and FRAP analysis

Cells were seeded in Lab-Tek chambered cover glass slides for microscopy. Expression of the opto-ZBP1^RHIM-only^ construct was induced with 1 µg/ml doxycycline for 24 h in the dark. For live-cell imaging), cells were exposed to a 2 min, 10V blue light pulse and imaged for 1h, with images acquired every 30 s. For snapshot images, cells were either kept in the dark or exposed to a 2 min, 10V blue light pulse, then fixed for 15 min with 4% formaldehyde and washed with PBS. Nuclei were stained using DAPI. For FRAP analysis, cells were either kept in the dark or imaged 3 h post-exposure to a 2min, 10V blue light pulse. FRAP images were acquired over a 5 min period at 5 s intervals. A region of interest was bleached by a 95% pulse of the 561 nm laser for 60 ms. All imaging was performed using a 63x/1.4 oil DIC objective (Plan-Apochromat, Zeiss) on a Zeiss Axio Observer.Z1 microscope driven by MetaMorph (MDS Analytical Technologies, Canada). The system was equipped with a Yokogawa spinning disk confocal head, an iLas multipoint FRAP module, an HQ2 CCD camera, a laser bench from Roper (405 nm 100 mW Vortran, 491 nm 50 mW Cobolt Calypso, and 561 nm 50 mW Cobolt Jive), and a stage-top incubator system (Live Cell Instruments) maintaining stable conditions at 37°C and 5% CO2.

### Image stream

Two million cells were seeded into 60 mm dishes 24h before start experiment. At endpoint, cells were detached using Trypsin/EDTA (0.05% Trypsin (Sigma-Aldrich, T4424); 0.032% EDTA (made in house)) and washed with PBS. Cells were stained with a live/dead stain (Invitrogen; eBioscience™ Fixable Viability Dye eFluor™ 780; 65-0865-14), to follow viability, and Hoechst (Thermofisher, H3570) to visualize the nuclei for 30 min at 4°C. Next, samples were washed in PBS and fixed, using the Foxp3/Transcription Factor Staining Buffer Set (eBioscience™; Invitrogen; 00-5523-00). Afterwards, cells were washed twice, resuspended in 50 µl PBS and stored at 4°C until flow cytometric analysis (Amnis Imagestream X MkII; Inspire). Quantification of ZBP1-eGFP signal was done using the IDEAS 6.3 software. Due to differences in the baseline expression of ZBP1 in polyclonal-sorted cell lines, the mask was adjusted accordingly to prevent detection of false positive events. A eGFP-positive aggregate was defined by following mask for clonal and polyclonal cell lines respectively; ‘range (peak (M02, CH02_GFP, bright 2) 0-20,0-1))’ and ‘range (peak (M02_GFP, bright 4.5),0-50,0-1))’. Quantification was done using the ‘Spot Count’ Feature on the GFP-aggregate mask.

### Cell death assay

Cell death measurements were done via repetitive imaging of cultured cells in the presence of cell-impermeable dye Sytox Green (Thermo Fisher, 10768273, 1/5000 dilution) or Propidium Iodide (Sigma-Aldrich, P-4170, 1/1000 dilution). Assays were done in a 96-well format, using Incucyte Zoom systems (Sartorius). Images were taken two hours and processed using the Incucyte Zoom software (Sartorius).

### Co-immunoprecipitation

Cells were washed and scraped in PBS and spun down for 5 minutes at 600g. Next, cells were lysed in 500 µl Amyloid Lysis Buffer (50 mM Tris pH 7.4; 137 mM NaCl; 1 mM EDTA; 1% Triton X100; 10% Glycerol; Protease inhibitor (cOmplete, sigma, 5056489001) for 15 minutes on turning wheel at 4°C. The samples were spun down for 10 minutes at 1000g. The supernatant was used to set up the immunoprecipitation (IP). A 50 µl samples was taken as input control. Magnetic flag beads (Sigma-Aldrich, M8823) or GFP-trap Magnetic particles M-270 (Chromotek, gtd-200) were washed 3 times with Amyloid Lysis buffer and then added to the samples. V5-IP was done with anti-V5 antibody (Invitrogen, R960-25), pre-conjugated with magnetic protein Dynabeads^TM^ (Thermofisher, 10003D) for 30 minutes at 4°C. IP was incubated for 3 hours (GFP trap/V5) or left overnight (flag). After 3 consecutive washes with Amyloid Lysis Buffer, the IP was resuspended in 50 µl of lysis buffer. Samples were further processed as described in ‘**Immunoblotting’**.

### Semi-Denaturing Detergent Gel Electrophoresis (SDD-AGE)

Transfected or infected cells were scraped in medium and spun down for 5 minutes at 1000g. After washing in PBS, cells were lysed in Amyloid Lysis Buffer (50 mM Tris pH 7.4; 137 mM NaCl; 1 mM EDTA; 1% Triton X100; 10% Glycerol; Protease inhibitor (cOmplete, sigma, 5056489001) for 30 minutes on ice. Samples were spun down for 10 minutes at 20 000g at 4°C. Afterwards, the supernatant was combined with loading buffer (2x TAE, 20% glycerol, 8% SDS, 0.08% Bromophenol blue) in a 4:1 ratio. The sample was incubated at room temperature for 10 minutes before loading it in an agarose gel (1% agarose; 0.1% SDS in TAE). The gel was run in TAE buffer, supplemented with 0.1% SDS at 60V for 2H30. A capillary transfer was performed on a PVDF membrane using TBS buffer (20mM Tris, ph7.4; 150 mM NaCl). Detecting protein on the membrane was done as described in **‘Immunoblotting’.**

### Immunoblotting

Lysates were made in Amyloid Lysis Buffer and lysed on ice for 30 minutes. Next, the sample was spun down for 10 minutes at 10 000g. Protein concentration was measured via the BCA protein assay kit (Pierce, ThermoFisher, 23225). Before loading samples on an acrylamide gel, samples were denatured with laemli buffer (250mM Tris, 10% SDS, 0.5% Bromophenol blue, 50% Glycerol) supplemented with 20 % β-Mercaptoethanol (Sigma-Aldrich, 441433A) in a 4:1 ratio and incubated at 95°C for 10 minutes. Proteins were loaded on 10% Tris-Acrylamide gels and separated by gel electrophoresis before they were transferred onto nitrocellulose membranes using semidry transfer systems (Hoefer^TM^). Membranes were blocked with 5% nonfat dry milk in TBS-T (Tris-buffered saline supplemented with 0.1% Tween 20) for 1 hour and then probed with primary antibody in 5% nonfat dry milk in TBS-T (see key resources table under tab ‘**Antibodies’**) overnight at 4°C. Membranes were washed in TBS-T and probed with horseradish peroxidase (HRP)–linked anti-mouse or anti-rabbit antibody in 5% nonfat dry milk or BSA (in TBS-T) for 1 hour at room temperature (information considering secondary antibodies; see table ‘**Antibodies’**). Blots were washed extensively before Protein visualisation using an enhanced chemiluminescence (ECL) reagent (Western Lightning Plus-ECL, PerkinElmer) on Amersham Imager 600 (General Electric).

### RNA isolation and RT-qPCR

RNA lysates were made with simplyRNA Tissue kit (Maxwell). Cells were incubated with a mix of 200 µl of lysis buffer and 200 µl isolation buffer, and subsequently loaded in the provided cartridges. The cartridge was prepared following manufacturer’s instructions, with 50 µl elution buffer and 10 µl DNaseI. Isolated RNA was directly used in the reconstitution assay or transcribed to cDNA using the SensiFast cDNA synthesis kit (Bioline, BIO-65054). Approximately 15 ng cDNA was used as input for quantitative Real-Time PCR (Lightcycler 480, Roche). SYBR-green based detection was done using SensiFast SYBR No-ROX kit (Bioline, BIO-98050). Expression data was normalised to B-Actin and Ywas using following formula: Rel. expression= (2^45-Ct(GOI)^)/(2^45-Ct(HKG)^). GOI: Gene Of Interest/ HKG: HouseKeeping Gene. For probe-based detection was done with TaqMan™ Gene Expression Master Mix (Thermofisher, 4369016). Expression data was normalysed to HPRT1 and ActB, using previous mentioned formula. qPCR primers and probes used in this study are listed in the table under the tab ‘**Primers**’.

### *In vitro* reconstitution assay

A T175 flask of ZBP1-GFP expressing cells at ∼90% confluency was detached with Trypsin/EDTA (0.05% Trypsin (Sigma-Aldrich, T4424); 0.032% EDTA (made in house)) and washed with PBS. Next, cells were resuspended in a hypotonic lysis buffer (10 mM Tris, pH 7.5; 5 mM KCl and 3mM MgCl2) supplemented with protease inhibitor (cOmplete, sigma, 5056489001), 200 µl per T175 flaks. Cells were lysed via mechanical disruption using a needle and syringe (30G, BD Micro-fine, 324826). Lysis of the cells was confirmed with Trypan blue (Merck, 11732). The sample was centrifuged for 5 minutes at 20 000g. Next, the supernatant containing protein extract was incubated with isolated RNA or DNA-polymers for 30 minutes at 37 °C before visualisation on TIRF microscope (Zeiss). In E3-competition assays, vaccinia virus protein E3 (Gentaur, CSB-EP322729VAA-50ug) was preincubated with the isolated RNA or GC-polymer for 30 minutes at 4°C. Concentrated ZBP1 lysate was then challenged with RNA/E3 or DNA/E3 mixture for 30 minutes at 37°C before visualisation on TIRF microscope.

### Statistical analysis

Statistical analyses were performed in Prism 8.3.0 (GraphPad Software). Statistical methods are described in the figure legends.

### Oligonucleotides

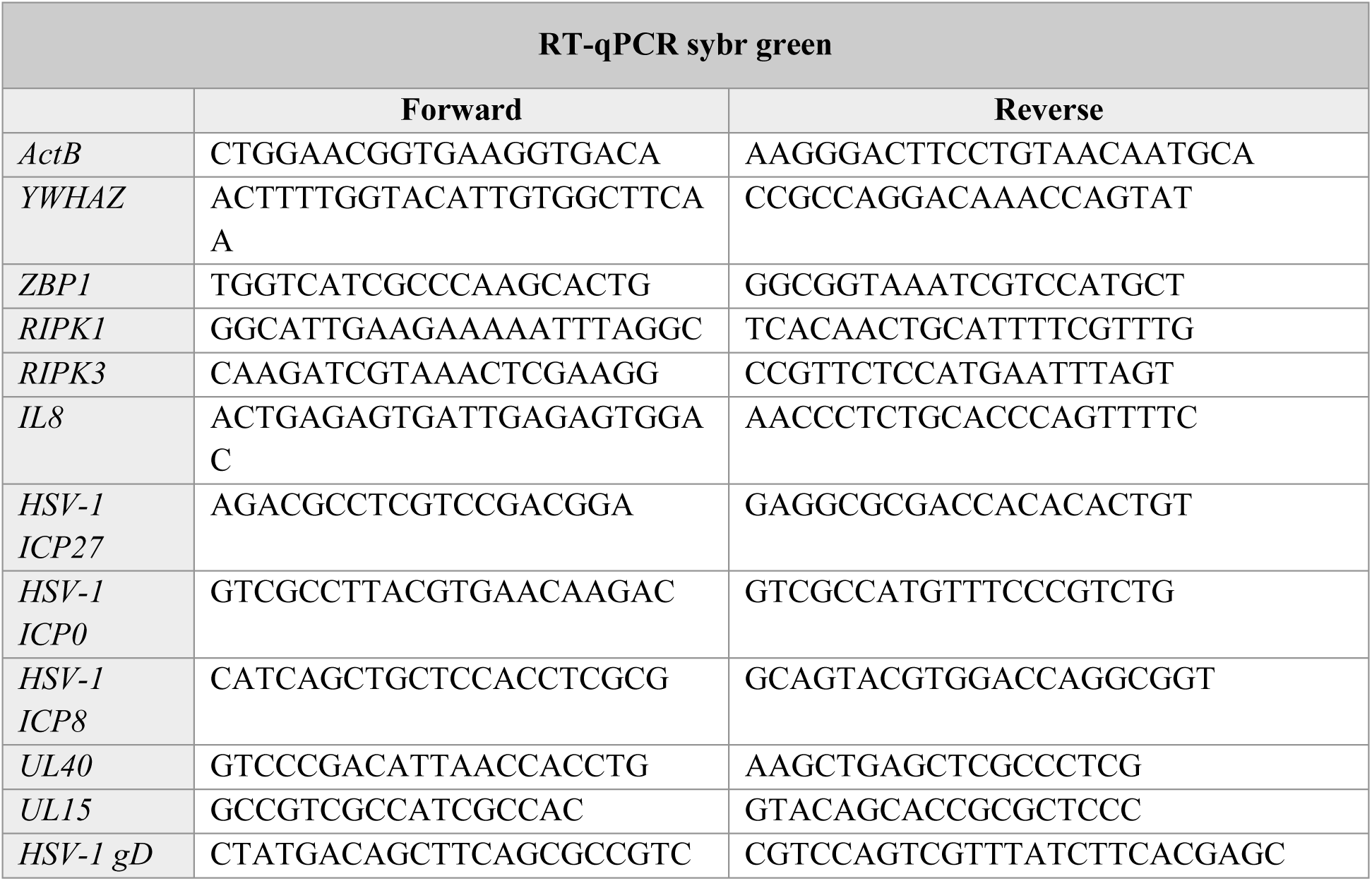
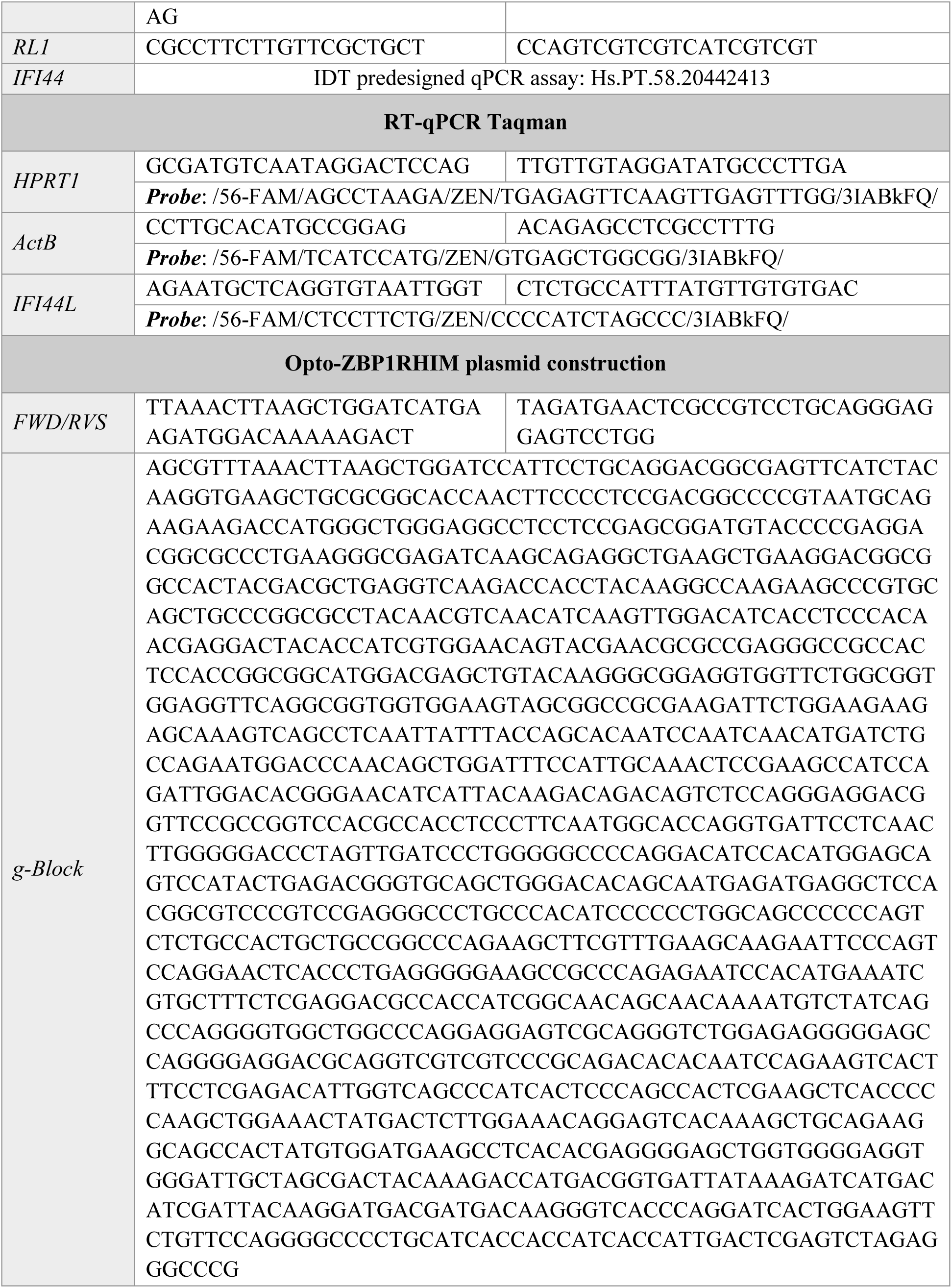

### Key resources table

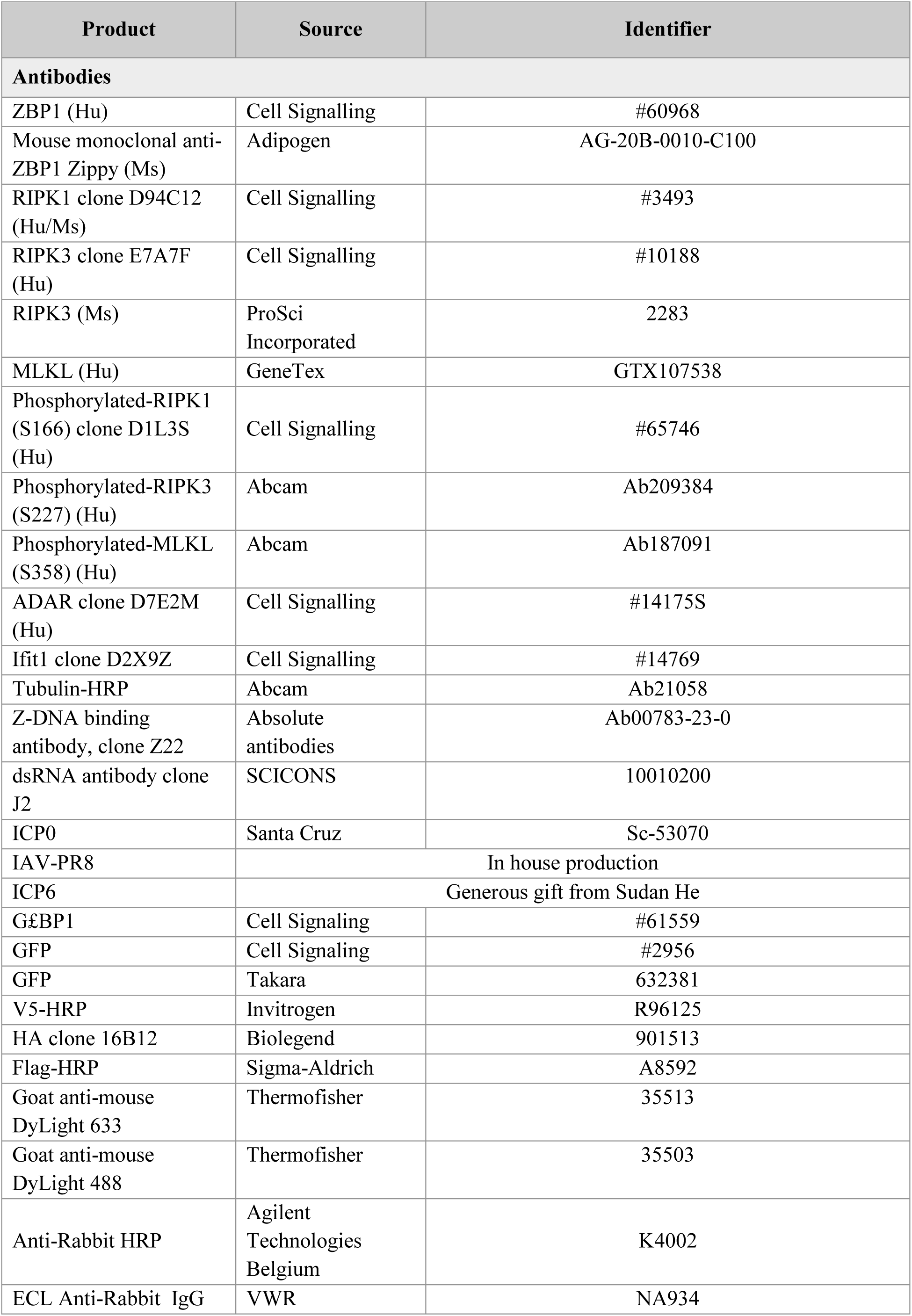
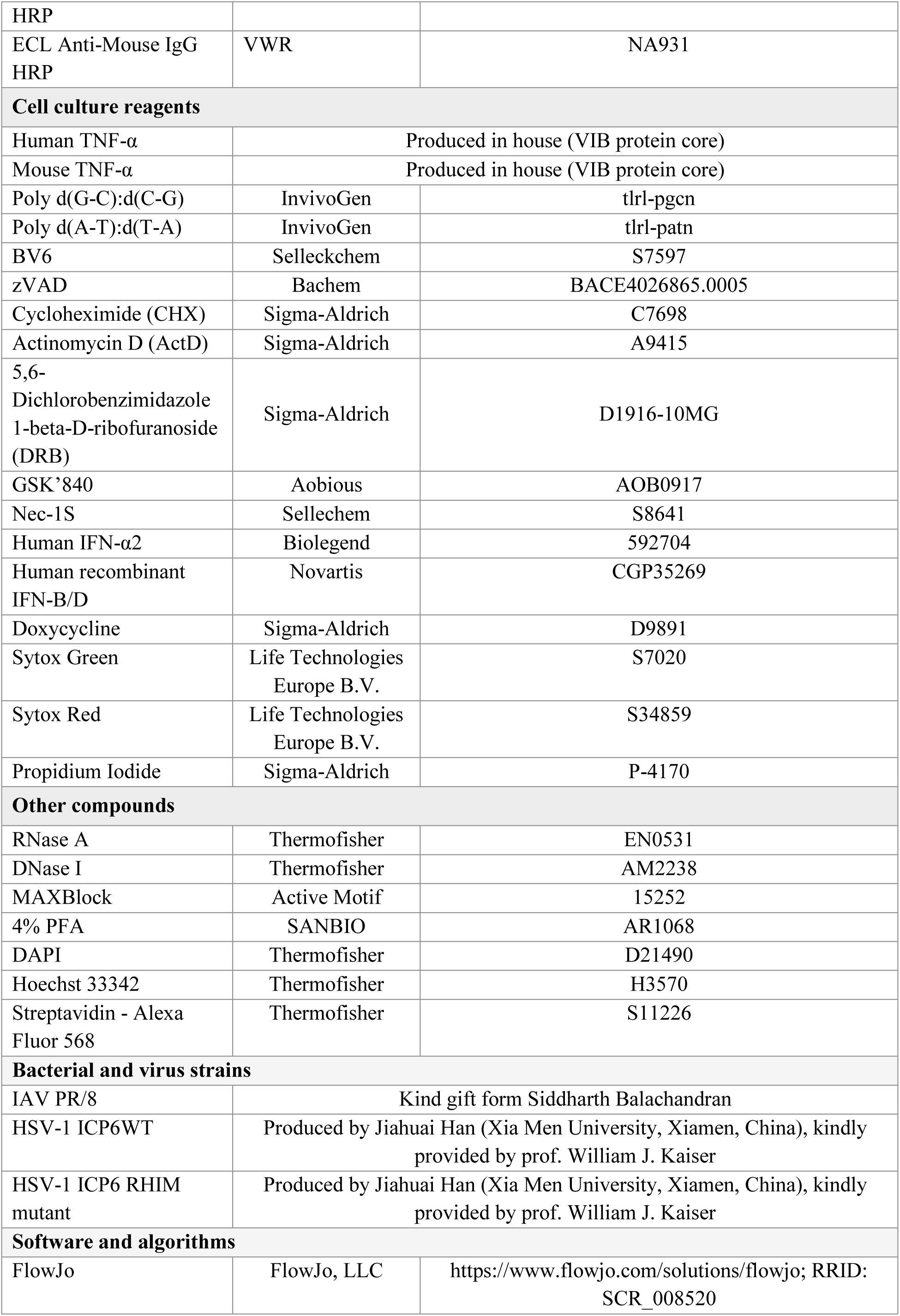
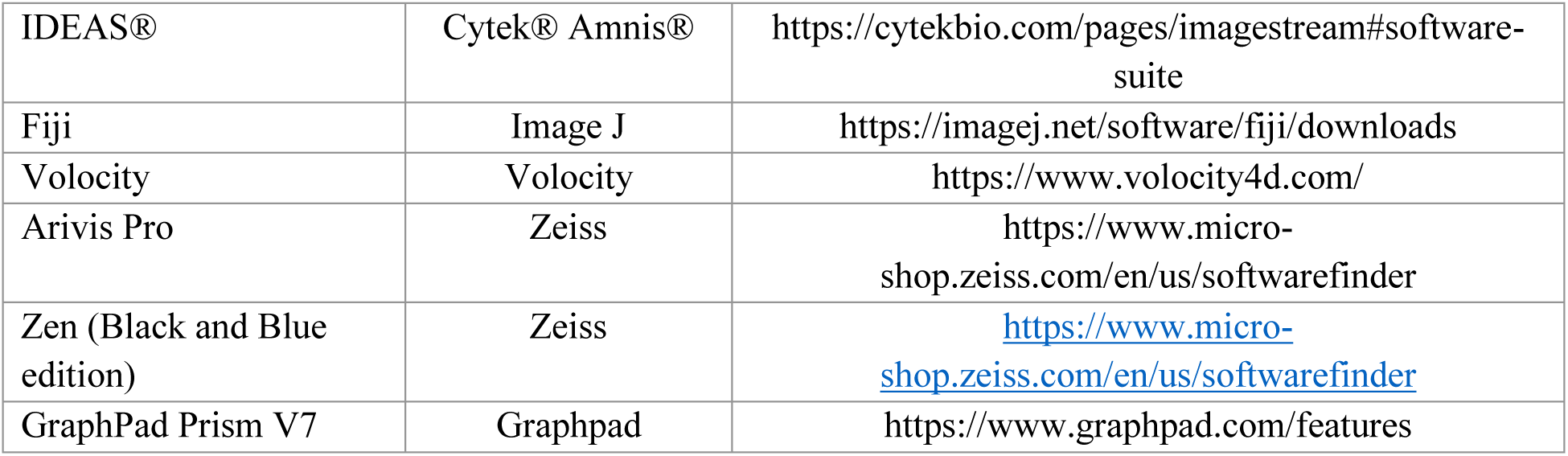

